# Glial Draper signaling triggers cross-neuron plasticity in bystander neurons after neuronal cell death

**DOI:** 10.1101/2023.04.09.536190

**Authors:** Yupu Wang, Ruiling Zhang, Sihao Huang, Parisa Tajalli-Tehrani Valverde, Meike Lobb-Rabe, James Ashley, Lalanti Venkatasubramanian, Robert A. Carrillo

**Affiliations:** Department of Molecular Genetics and Cellular Biology, University of Chicago, Chicago, IL 60637; Neuroscience Institute, University of Chicago, Chicago, IL 60637; Committee on Development, Regeneration, and Stem Cell Biology, University of Chicago, Chicago, IL 60637; Program in Biochemistry and Molecular Biophysics, University of Chicago, Chicago, IL 60637; Program in Cell and Molecular Biology, University of Chicago, Chicago, IL 60637; Department of Zoology, University of Cambridge, United Kingdom; Howard Hughes Medical Institute, Janelia Research Campus, Ashburn, VA 20147

## Abstract

Neuronal cell death and subsequent brain dysfunction are hallmarks of aging and neurodegeneration, but how the nearby healthy neurons (bystanders) respond to the cell death of their neighbors is not fully understood. In the *Drosophila* larval neuromuscular system, bystander motor neurons can structurally and functionally compensate for the loss of their neighbors by increasing their axon terminal size and activity. We termed this compensation as cross-neuron plasticity, and in this study, we demonstrated that the *Drosophila* engulfment receptor, Draper, and the associated kinase, Shark, are required in glial cells. Surprisingly, overexpression of the Draper-I isoform boosts cross-neuron plasticity, implying that the strength of plasticity correlates with Draper signaling. Synaptic plasticity normally declines as animals age, but in our system, functional cross-neuron plasticity can be induced at different time points, whereas structural cross-neuron plasticity can only be induced at early stages. Our work uncovers a novel role for glial Draper signaling in cross-neuron plasticity that may enhance nervous system function during neurodegeneration and provides insights into how healthy bystander neurons respond to the loss of their neighboring neurons.

## Introduction

One of the most remarkable features of the brain is its plasticity – the capacity of neural circuits and synapses to adapt to experiences or perturbations by modifying their activity and even their morphology. Many forms of synaptic plasticity have been reported (Citri and Malenka, 2008), such as long-term potentiation (LTP) (Bliss and Lømo, 1973), long-term depression (LTD) (Mulkey and Malenka, 1992), Hebbian plasticity (Hebb, 1949), presynaptic homeostatic potentiation (PHP) (Burrone et al., 2002; Murthy et al., 2001), presynaptic homeostatic depression (PHD) (Daniels et al., 2004), and others. These mechanisms allow individual synapses to alter their composition and activity during development, learning, injury, and disease. However, during aging and neurodegeneration when substantial neuronal cell death occurs, these plasticity paradigms cannot alleviate the functional defects since the synaptic connections are disrupted, and in many cases, the synapses no longer exist (Mattson and Magnus, 2006; Peters et al., 2008). Interestingly, several studies indicate that neuronal injury or death may alter the structural and functional properties of nearby healthy “bystander” neurons (Aponte-Santiago et al., 2020; Hsu et al., 2020; Wang et al., 2021). This model highlights that bystander neurons may be an overlooked resource to compensate the nervous system defects during neuronal dysfunction and death.

The first report (to our knowledge) that healthy bystander neurons can respond to injury or death of neighboring neurons was nearly a century ago at the vertebrate neuromuscular junction (NMJ) – denervation of a muscle fiber can induce sprouting of nearby motor neurons (MNs) and eventually restore the circuit function (Brown et al., 1981; Edds, 1949; Edds, 1953; Hoffman, 1950). A similar phenotype was observed at invertebrate NMJs. In crustaceans, killing the common MN that innervates multiple muscles induced the bystander MN that co-innervates the muscle to increase its NMJ size and quantal content (Parnas et al., 1984). In leech embryos, physical removal of the S interneuron that resides in one ganglion led to compensation from an S interneuron in a nearby ganglion. The healthy S interneuron extended axons into the lesioned ganglion and restored the synaptic function (Modney and Muller, 1994). Recent studies at the *Drosophila* larval NMJ reported similar observations using a genetic approach to ablate MNs (Aponte-Santiago et al., 2020; Han et al., 2022; Wang et al., 2021). Most *Drosophila* larval muscles are innervated by two excitatory glutamatergic MNs, known as type-I big MN (Ib MN) and type-I small MN (Is MN) based on their terminal bouton size (Choi et al., 2004; Lnenicka and Keshishian, 2000). In each hemisegment, ∼29 Ib MNs innervate thirty muscle fibers mostly in a one-to-one manner (Hoang and Chiba, 2001; Wang et al., 2022), whereas two Is MNs innervate separate groups of muscles and therefore, are known as the common exciters (Takizawa et al., 2007). Genetic ablation of Is MNs led to an expanded NMJ and an elevated excitatory post-synaptic potential (EPSP) amplitude in healthy bystander Ib MNs (Aponte-Santiago et al., 2020; Han et al., 2022; Wang et al., 2021). Four Ib MNs have been examined and three of them showed compensation at differing levels upon Is MN ablation (MN1-Ib, MN4-Ib and MN6-Ib). In addition, ablating Ib MNs (MN1-Ib) did not induce compensation in the corresponding Is MN, suggesting MNs differ in their ability to compensate (Aponte-Santiago et al., 2020). These vertebrate and invertebrate studies provide strong support for healthy bystander neurons in restoring synaptic function upon the loss of their neighbors. However, the mechanisms by which these bystander neurons respond to the death of their neighbors are largely unknown. In this study, we referred to plasticity changes induced by neuronal cell death as “cross-neuron plasticity” and explored the underlying mechanisms.

In the cross-neuron plasticity studies described above, the dying neuron and the bystander neurons do not physically contact or synaptically connect with each other, suggesting a third party is likely involved in sensing the injury and spreading the signal. Glial cells are an attractive candidate because of their close association with neurons and essential roles in neural development, synaptic plasticity, and injury responses (Freeman, 2015; Yildirim et al., 2019). During development and metamorphosis when substantial axon or dendrite pruning occurs, and during injury induced neuronal degeneration, glial cells detect and engulf neuronal debris through a conserved signaling pathway mediated by an engulfment receptor, MEGF10 and Jedi (vertebrates, (Lee et al., 2020; Wu et al., 2009))/CED-1 (C-elegans, (Zhou et al., 2001))/Draper (Drosophila, (Awasaki et al., 2006; Hoopfer et al., 2006; MacDonald et al., 2006)). Draper contains an ITAM domain found in many mammalian immunoreceptors which can be phosphorylated by Src42a to allow binding of an SH2 domain kinase, Shark (Ziegenfuss et al., 2008). The Draper/Shark complex recruits the glial membrane for engulfment through dCed-6 and Rac1 and activates engulfment gene expression through the dJNK pathway (Lu et al., 2014; MacDonald et al., 2013; Ziegenfuss et al., 2012). The necessity of the Draper/Shark pathway in glia-mediated clearance of neuronal debris following injury led to an appealing hypothesis that this engulfment pathway may serve as the trigger to initiate cross-neuron plasticity. Importantly, a recent study found that severing of sensory axons in the adult wing influenced cargo transport in bystander axons in a Draper-dependent mechanism (Hsu et al., 2020). However, whether and how the morphology and physiology of bystander neurons are affected is not known.

In this study, we utilized the larval neuromuscular system to examine how bystander neurons detect loss of neighboring neurons. First, we genetically ablated Is MNs and found that Draper is required for clearance of the Is MN axon and cell body debris. Next, we examined structural and functional cross-neuron plasticity in healthy bystander Ib MNs and found that the Draper/Shark signaling pathway is required in glial cells. In addition, Draper-I is the specific isoform mediating cross-neuron plasticity and elevating Draper-I expression can boost plasticity. These data provide insights about how cross-neuron plasticity is detected in the neuromuscular system. To further explore cross-neuron plasticity, we performed age-dependent Is MN ablation and found that Ib MN functional, but not structural, plasticity can be induced at all larval stages. Finally, to determine the behavioral consequences of Ib cross-neuron plasticity, we examined larval locomotion and observed elevated crawling speed. Overall, these data support an important role for healthy bystander neurons in detecting and responding to the nearby neuronal loss.

## Results

### Draper is required for debris clearance after Is MN ablation

The *Drosophila* engulfment receptor, Draper, is implicated in axonal debris clearance in several *Drosophila* nerve injury models including axotomy of olfactory and wing sensory neurons (Hsu et al., 2020; MacDonald et al., 2006; Purice et al., 2016). However, whether Draper is required for the clearance of axonal debris generated by programmed cell death and the subsequent cross-neuron plasticity is not clear. Here, we genetically ablated Is MNs by ectopic expression of the cell death genes head involution defective (*hid*) and reaper (*rpr*) in a *draper* mutant background (*draper^Δ5^*) and examined Is MN debris clearance. We used a Is MN specific GAL4, *A8-GAL4* (hereafter named *Is-GAL4*), to label and ablate Is MNs. In a prior study, we demonstrated that expression of *Is-GAL4* begins at embryonic stage 14 and efficiently ablated Is MNs by early first instar stage (Wang et al., 2021). In *Is>GFP* larvae, GFP labels the Is MN axons (Figure 1A and A’), and co-expression of *rpr,hid* (*Is>GFP,rpr,hid*) lead to ablation of Is MNs and complete removal of the Is MN axons (Figure 1B and B’). However, ablation of Is MNs in *draper* mutant animals lead to significant accumulation of GFP in the segmental nerve that co-localized with the glial cell marker, Repo (Figure 1C-C’, D-D’ and E), suggesting a failure of axonal debris clearance (Yildirim et al., 2019). Next, we examined the ventral nerve cord (VNC) where Is MN cell bodies and dendrites are located. We found complete clearance of Is MN debris in the VNC in *Is>GFP,rpr,hid* first instar larvae (Figure 1F,G); however, ablation in a *draper* mutant background resulted in significant GFP retention (Figure H-J), similar to the accumulation at segmental nerve. In summary, we showed that Draper is required to efficiently remove neuronal debris induced by programmed cell death, in both the segmental nerve and the VNC.

**Figure 1.**
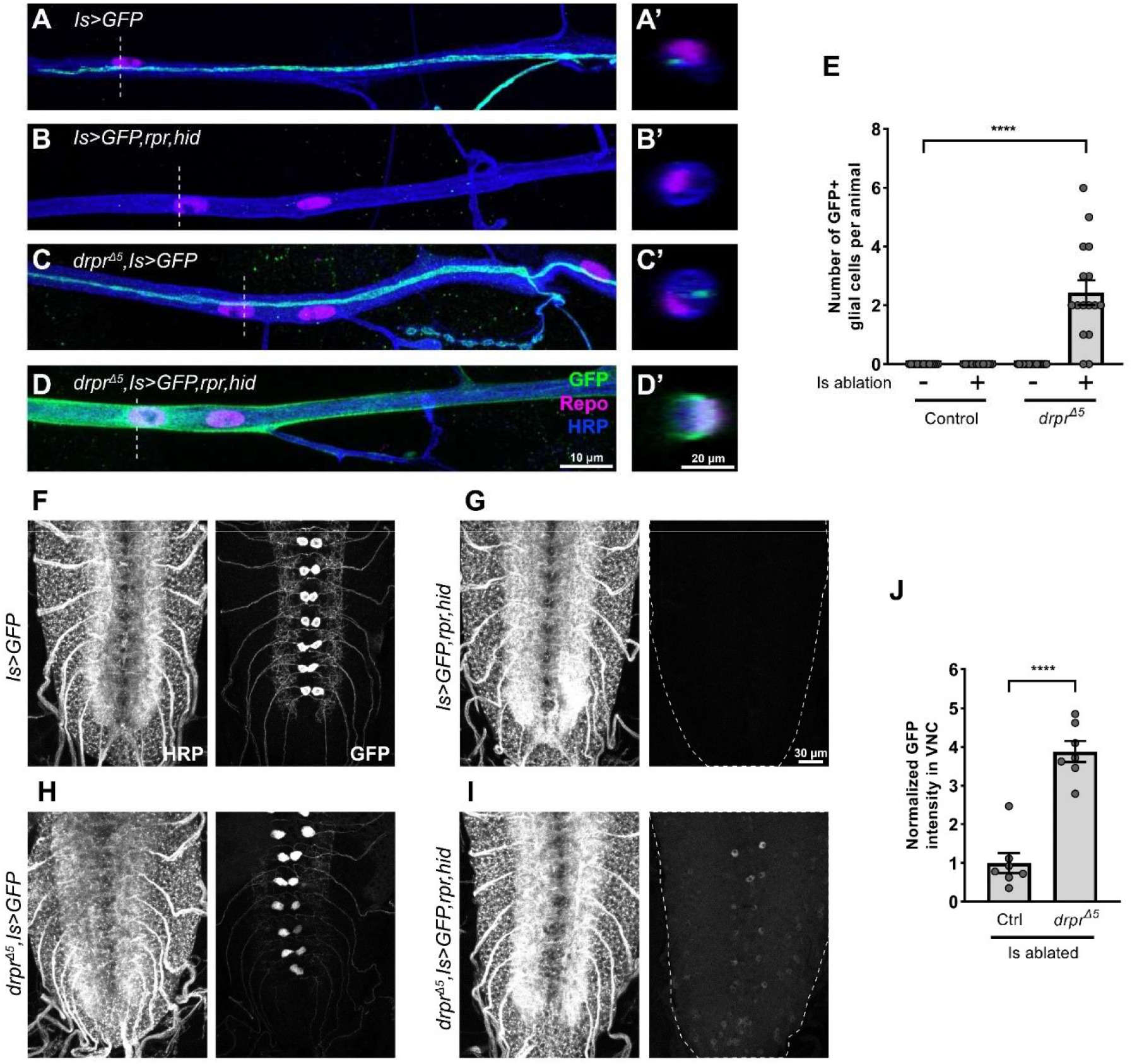
Draper is required for debris clearance after Is MN ablation. A-D and A’-D’. Axon bundles in third instar (A) no ablation control (*Is>GFP*), (B) Is ablated (*Is>GFP,hid,rpr*), (C) *draper* mutant with no ablation (*drpr^Δ5^, Is>GFP*) and (D) *draper* mutant with Is ablated (*drpr^Δ5^, Is>GFP,hid,rpr*) larvae, labeled with GFP (green), Repo (glial cell marker, magenta) and HRP (neuronal marker, blue). Grey dash lines indicate the position of cross sections (A’-D’). Significant GFP positive debris accumulated in glial cells when ablating Is MNs in a *draper* mutant background (D and D’). E. Quantification of the number of GFP+ glial cells per animal. F(3,57)=14.09, p<0.0001, one-way ANOVA. N (larvae) =15, 14, 16, 16. F-I. VNCs of first instar larvae of displayed genotypes labeled with HRP (left panel) and GFP (right panel). Note the significant amount of GFP+ signal remaining in (I) after removing *draper*. J. Quantification of GFP intensity in VNC. t(12)=7.703, p<0.0001, unpaired t-test. N (VNCs) =7, 7. Error bars indicate ± SEM, ****p<0.0001.

### Draper is required for cross-neuron plasticity

We reasoned that clearance of the MN debris may be part of the signaling pathway that initiates cross-neuron plasticity. To test this hypothesis, we examined the role of Draper as it was required for efficiently removing MN debris upon cell ablation. We genetically ablated Is MNs and examined a specific bystander Ib MN that innervates the dorsal muscle 4 (MN4-Ib), because in a previous study, this MN displayed robust structural and functional plasticity when the adjacent Is MN was ablated (Wang et al., 2021). We first examined the NMJ size of the MN4-Ib to determine structural plasticity. Ablating Is MNs in a wild type background led to an increase of MN4-Ib bouton number, as previously reported (Wang et al., 2021) (Figure 2A, B and E). Interestingly, this NMJ expansion is not observed when Is MNs are ablated in a *draper* mutant background (Figure 2C-E), suggesting Draper is required for structural plasticity.

**Figure 2.**
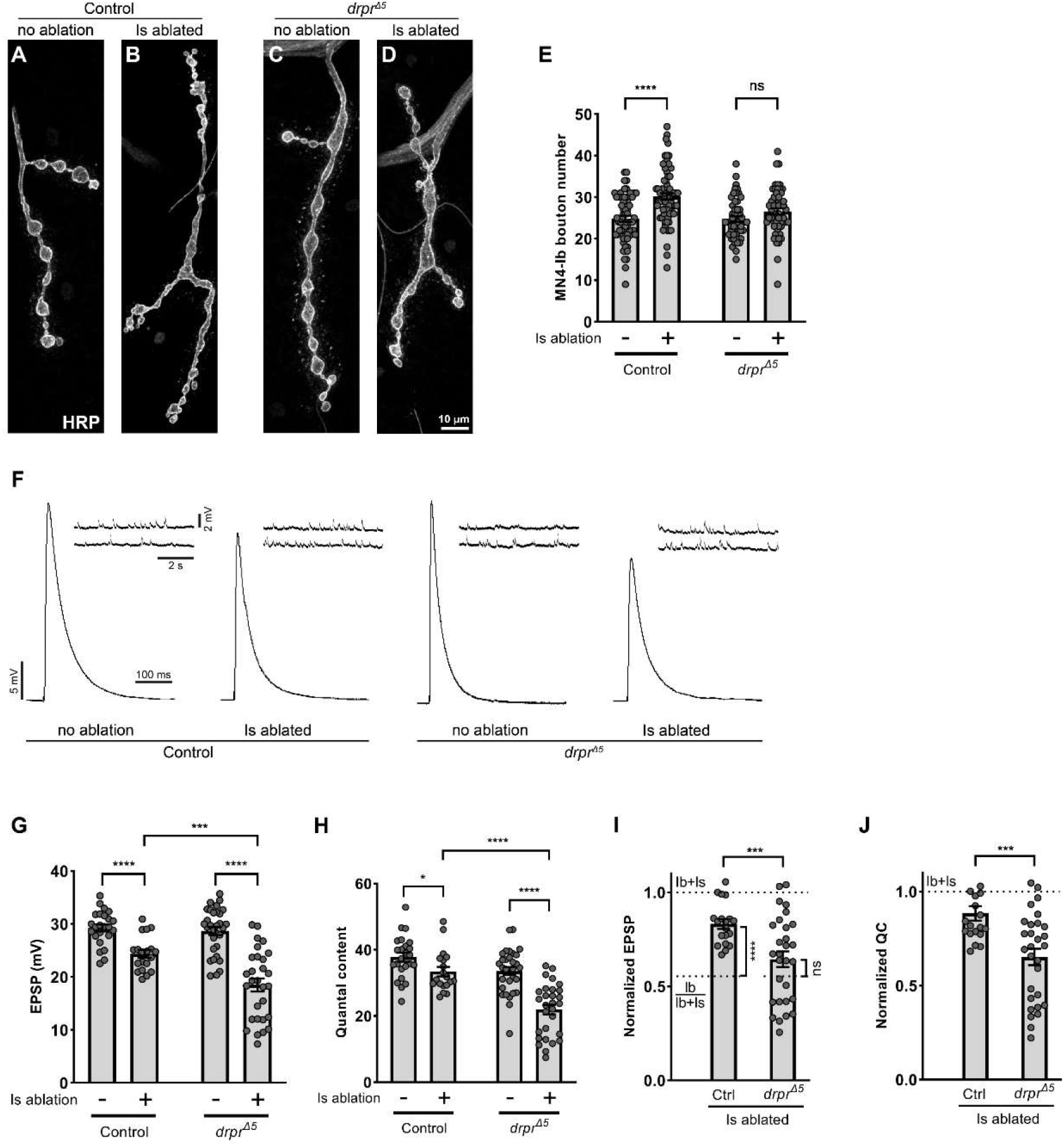
Draper is required for cross-neuron plasticity. A-D. NMJs of MN4-Ib in third instar (A) no ablation control (*Is>GFP*), (B) Is ablated (*Is>GFP,hid,rpr*), (C) *draper* mutant with no ablation (*drpr^Δ5^, Is>GFP*) and (D) *draper* mutant with Is ablated (*drpr^Δ5^, Is>GFP,hid,rpr*) larvae, labeled with HRP (grey). The NMJ was expanded in control Is ablated larvae due to cross-neuron plasticity (B), and this expansion is blocked in a *draper* mutant background (D). E. Quantification of MN4-Ib bouton numbers in no ablation and Is ablated larvae in control and *drpr^Δ5^* backgrounds. Control (n = 66 and 69 NMJs), t(133)=5.030, p<0.0001, unpaired t-test. *drpr^Δ5^* (N = 53 and 56 NMJs), t(107)=1.838, p=0.0688, unpaired t-test. F. EPSP and mEPSP traces from no ablation and Is ablated larvae in control and *drpr^Δ5^* backgrounds. G. Quantification of EPSP amplitude of no ablation and Is ablated larvae in control and *drpr^Δ5^* backgrounds. Control, t(41)=4.924, p<0.0001, unpaired t-test. *drpr^Δ5^*, t(48.04)=7.011, p<0.0001, unpaired t-test with Welch’s correction. Is ablated control vs Is ablated in *drpr^Δ5^*, t(43.54)=4.075, p=0.0002, unpaired t-test with Welch’s correction. H. Quantification of quantal content of no ablation and Is ablated larvae in control and *drpr^Δ5^* backgrounds. Control, t(41)=2.224, p=0.0317, unpaired t-test. *drpr^Δ5^*, t(58)=6.194, p<0.0001, unpaired t-test. Is ablated control vs Is ablated in *drpr^Δ5^*, t(46)=5.304, p<0.0001, unpaired t-test. I. Quantification of normalized EPSP of Is ablated larvae in control and *drpr^Δ5^* backgrounds. Is ablated control vs *drpr^Δ5^*, t(43.20)=3.753, p=0.0005, unpaired t-test with Welch’s correction. Is ablated control vs Ib/Ib+Is, t(29)=6.506, p<0.0001, unpaired t-test. Is ablated in *drpr^Δ5^* vs Ib/Ib+Is, t(37.82)=1.524, p=0.1357, unpaired t-test with Welch’s correction. J. Quantification of normalized quantal content of Is ablated larvae in control and *drpr^Δ5^* backgrounds. t(46)=3.730, p=0.0005, unpaired t-test. For G-I, N (NMJs) = 24, 19, 31, 29. Error bars indicate ± SEM, ns = non-significant, *p<0.05, ***p<0.001, ****p<0.0001.

Next, we tested the role of Draper in functional cross-neuron plasticity. Muscle 4 receives innervation from both MN4-Ib and the dorsal Is MN, and these neurons are normally activated simultaneously during electrophysiology recordings (Figure S1A). However, to understand the changes of Ib MN activity before and after ablation, we needed to isolate Ib MN activity from a wild type animal where both Ib and Is MNs are present. One approach to separate the activity of these MNs is by tuning the stimulating voltage, as Ib MNs have a lower stimulating threshold than Is MNs (Lnenicka and Keshishian, 2000; Schaefer et al., 2010). Using GCaMP imaging to visualize the activated MN together with electrophysiology recording from the muscle (Figure S1A), we recorded Ib and Ib+Is excitatory post-synaptic potentials (EPSPs) (Wang et al., 2021). We recorded a smaller EPSP which is generated by stimulation of the Ib MN alone, and a larger EPSP which is generated by activation of both Ib and Is MNs (Figure S1B). We normalized the Ib EPSP to Ib+Is EPSP and found that MN4-Ib contributes approximately 56% to the total EPSP, similar to our previous observation (Wang et al., 2021). We will therefore use this ratio (Ib/Ib+Is) to indicate the MN4-Ib baseline activity. Next, we recorded spontaneous and evoked activity in wild type and *draper* mutant animals, with or without Is MN ablation (Figure 2F). We did not observe any significant changes with spontaneous release in the *draper* mutant background as measured by frequency or amplitude of the spontaneous EPSP (also known as miniature EPSP, mEPSP) (Figure 2F and Figure S2). Examination of evoked activity revealed significantly smaller Ib EPSPs and quantal content when ablating Is MNs in a *draper* mutant background compared to control Is ablated animals (Figure 2G and 2H). To better illustrate the data, we normalized the Is ablated EPSP and quantal content to respective controls (i.e. control Is ablated EPSP is normalized to control non-ablated EPSP), and compared the normalized data across genotypes, together with the Ib baseline activity (Ib/Ib+Is). We found that upon Is MN ablation, the Ib EPSP is significantly higher than Ib baseline activity, suggesting a robust functional cross-neuron plasticity (Figure 2I, comparing Ctrl to Ib/Ib+Is). However, this compensation is absent in *draper* mutant animals (Figure 2I, comparing *drpr^Δ5^* to Ib/Ib+Is). In addition, direct comparison of the normalized EPSP and quantal content confirmed a loss of functional cross-neuron plasticity in the mutant background (Figure 2I and 2J). Taken these together, our data suggest that Draper is required for both structural and functional cross-neuron plasticity.

### Draper is required in glial cells to mediate cross-neuron plasticity

Draper is expressed in multiple cell types, including glial cells and muscles (Figure S3A-B), where it functions as an engulfment receptor (Fuentes-Medel et al., 2009; Logan et al., 2012). MNs interact extensively with glial cells in both the VNC and segmental nerve bundles as well as with muscles at the NMJ. Therefore, we sought to determine where Draper function is required for cross-neuron plasticity. Here, we first validated a *draper* RNAi line and found that expressing RNAi using the glial cell driver, *Repo-GAL4*, or the muscle driver, *Mef2-GAL4*, eliminated Draper expression in respective cells (Figure S3C-E).

We then examined cross-neuron plasticity in animals with cell specific *draper* knockdown. Ablating Is MNs in controls lead to an increase of Ib bouton number (Figure 3A,B,I), and this NMJ expansion was blocked by *draper* double knockdown in both glial cells and muscles (Figure 3C,D,I), similar to the *draper* mutant phenotype (Figure 2E). *draper* single knockdown in glial cells blocked the elevation of Ib bouton numbers (Figure 3E,F,I) while the muscle knockdown did not (Figure 3G,H,I), suggesting that Draper is specifically required in glial cells. Next, we performed electrophysiology analyses in these knockdown conditions (Figure S4). We found that knocking down *draper* in glial cells fully blocks the Ib EPSP compensation, when comparing the normalized EPSP to control Is ablated animals or to the Ib baseline activity (Figure 3J). Similarly, the elevated Ib quantal content induced by Is ablation is blocked when knocking down *draper* in glial cells (Figure 3K). Removing *draper* in muscles also caused a decrease of Ib EPSP compensation (Figure 3J), but the quantal content is not significantly different than control Is ablated animals (Figure 3K), suggesting Draper is only partially required in muscles for cross-neuron plasticity. Finally, *draper* knockdown in both glial cells and muscles fully blocked functional plasticity, similar to knockdown in glial cells alone (Figure 3J and 3K). Taken together, we showed that Draper is required in the glial cells to mediate cross-neuron plasticity.

**Figure 3.**
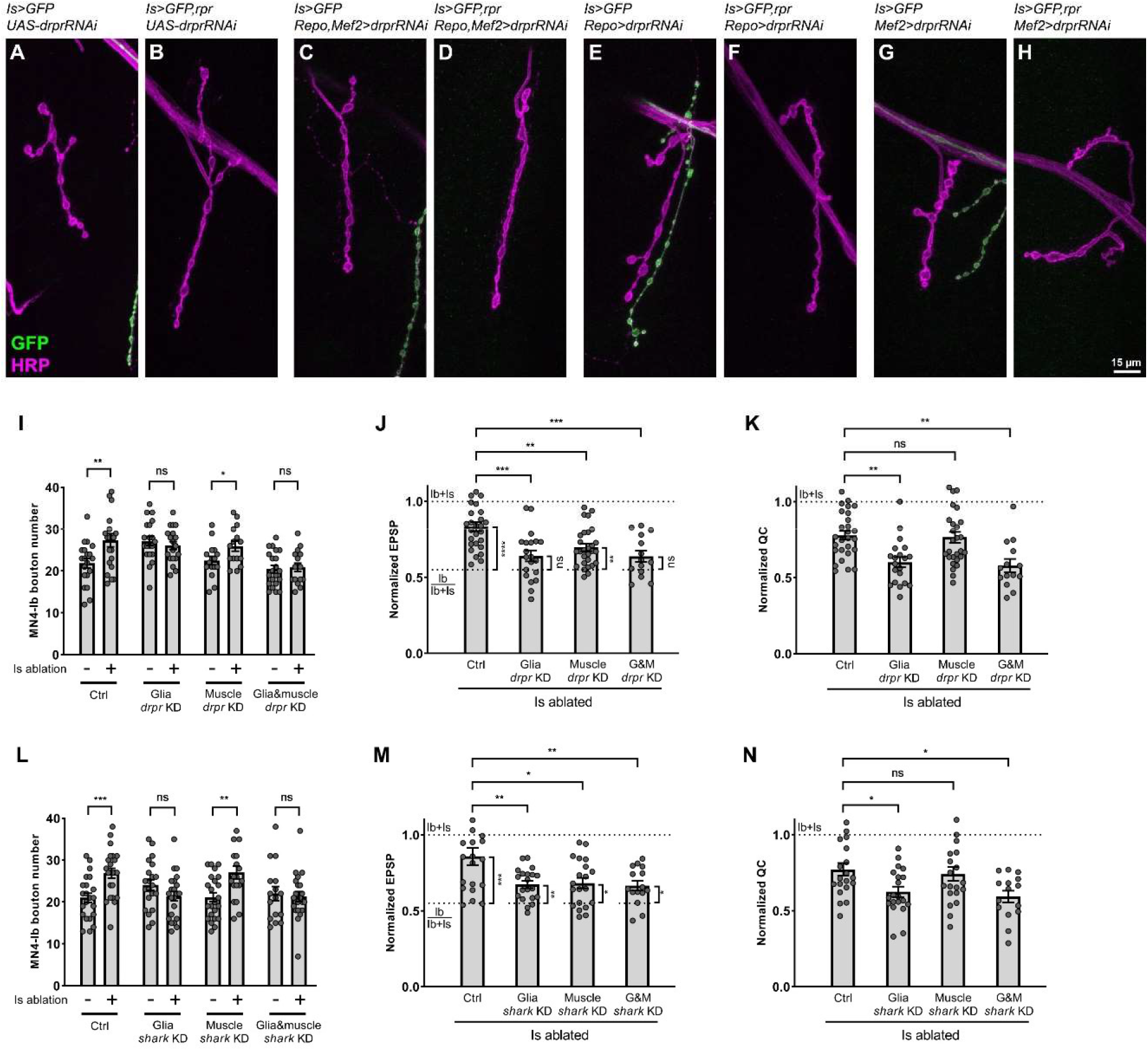
Draper and Shark are required in glial cells for cross-neuron plasticity. A-H. NMJs of MN4-Ib in third instar no ablation and Is ablated larvae in control, glia *draper* knockdown, muscle *draper* knockdown, and double knockdown backgrounds, labeled with GFP (green) and HRP (magenta). NMJ expansion was observed upon Is MN ablation (B), and this expansion is absent in glia *draper* knockdown (D) or double knockdown (H) backgrounds. I. Quantification of MN4-Ib bouton number between no ablation and Is ablated larvae in control, glia *draper* knockdown, muscle *draper* knockdown, and double knockdown backgrounds. Control (N = 20 and 21 NMJs), t(39)=2.822, p=0.0075, unpaired t-test. Glia *draper* knockdown (N = 20 and 19 NMJs), t(37)=0.7525, p=0.4565, unpaired t-test. Muscle *draper* knockdown (N = 16 and 15 NMJs), t(29)=2.204, p=0.0356, unpaired t- test. Double knockdown (N = 21 and 16 NMJs), t(35)=0.2965, p=0.7686, unpaired t-test. J. Quantification of normalized EPSP of Is ablated larvae in control, glia *draper* knockdown, muscle *draper* knockdown, and double knockdown backgrounds. F(3, 85)=9.191, p<0.0001, one-way ANOVA. Is ablated control vs glia *draper* knockdown, p=0.0001. Is ablated control vs muscle *draper* knockdown, p=0.0042. Is ablated control vs double knockdown, p=0.0006. Is ablated control vs Ib/Ib+Is, t(37)=5.462, p<0.0001, unpaired t-test. Is ablated in glia *draper* knockdown vs Ib/Ib+Is, t(30)=1.483, p=0.1486, unpaired t-test. Is ablated in muslce *draper*knock down vs Ib/Ib+Is, t(38)=3.178, p=0.0029, unpaired t-test. Is ablated in double knockdown vs Ib/Ib+Is, t(24)=1.540, p=0.1367, unpaired t-test. K. Quantification of normalized quantal content of Is ablated larvae in control, glia *draper* knockdown, muscle *draper* knockdown, and double knockdown backgrounds. F(3, 85)=8.263, p<0.0001, one-way ANOVA. Is ablated control vs glia *draper* knockdown, p=0.0028. Is ablated control vs muscle *draper*knock down, p=0.9913. Is ablated control vs double knockdown, p=0.0052. For J and K, N (NMJs) = 27, 20, 28, 14. L. Quantification of MN4-Ib bouton number in no ablation and Is ablated larvae in control, glial *shark* knockdown, muscle *shark* knockdown, and double knockdown backgrounds. Control (N = 22 and 23 NMJs), t(43)=3.598, p=0.0008, unpaired t-test. Glial *shark* knockdown (N = 21 and 22 NMJs), t(41)=1.566, p=0.1250, unpaired t-test. Muscle *shark* knockdown (N = 23 and 19 NMJs), t(40)=3.220, p=0.0025, unpaired t-test. Double *shark* knockdown (N = 16 and 22 NMJs), t(36)=0.3390, p=0.7366, unpaired t- test. M. Quantification of normalized EPSP of Is ablated larvae in control, glia *shark* knockdown, muscle *shark* knockdown, and double knockdown backgrounds. F(3, 71)=5.533, p=0.0018, one-way ANOVA. Is ablated control vs glia *shark* knockdown, p=0.0062. Is ablated control vs muscle *shark* knockdown, p=0.0105. Is ablated control vs double knockdown, p=0.0093. Is ablated control vs Ib/Ib+Is, t(28.04)=4.485, p=0.0001, unpaired t-test with Welch’s correction. Is ablated in glia *shark* knockdown vs Ib/Ib+Is, t(30)=2.798, p=0.0089, unpaired t-test. Is ablated in muscle *shark* knockdown vs Ib/Ib+Is, t(30)=2.329, p=0.0268, unpaired t-test. Is ablated in double knockdown vs Ib/Ib+Is, t(25)=2.254, p=0.0332, unpaired t-test. N. Quantification of normalized quantal content of Is ablated larvae in control, glia *shark* knockdown, muscle *shark* knockdown, and double knockdown backgrounds. F(3, 71)=4.437, p=0.0065, one-way ANOVA. Is ablated control vs glia *shark* knockdown, p=0.0466. Is ablated control vs muscle *shark* knockdown, p=0.9470. Is ablated control vs double knockdown, p=0.0214. For M and N, N (NMJs) = 20, 20, 20, 15. Error bars indicate ± SEM, ns = non-significant, *p<0.05, **p<0.01, ***p<0.001.

### The Draper co-factor, Shark, is required in cross-neuron plasticity

Previous studies revealed that Draper acts together with an essential kinase, Shark, to regulate engulfment and target gene expression (Doherty et al., 2014; Lu et al., 2017; Ziegenfuss et al., 2008). Therefore, we hypothesized that Shark may also be required for cross-neuron plasticity. We took an RNAi approach to specifically knock down *shark* in glial cells and muscles. We first assayed for structural NMJ changes (Figure S5) and observed that *shark* knockdown in glial cells blocks the Ib structural compensation induced by Is ablation, whereas knockdown in muscles does not (Figure 3L). Additionally, *shark* knockdown in both glial cells and muscles also blocked Ib structural compensation, similar to glial cell knockdown, suggesting that Shark is indeed required in glial cells for structural plasticity (Figure 3L). Next, we measured the Ib functional compensation in *shark* knockdown animals (Figure S6). We found that knocking down *shark* in glial cells leads to a significant loss of Ib functional plasticity, as the EPSP and quantal content are both decreased compared to controls (Figure 3M and 3N). Muscle knockdown led to a slight decrease of the EPSP, but no significant change of the normalized quantal content (Figure 3M and 3N), suggesting that muscles may have a limited role, as we previously observed in *draper* knockdown experiments (Figure 3J and 3K). In addition, simultaneous knockdown of *shark* in both muscles and glia mimicked glial knockdown, further confirming a significant role of *shark* in glial cells in cross-neuron plasticity (Figure 3M and 3N). In summary, our data suggested that Draper and its co-factor, Shark, are both required for cross-neuron plasticity.

### Overexpression of Draper-I boosts cross-neuron plasticity

After demonstrating that Draper is required for cross-neuron plasticity, we wondered if we could boost plasticity by overexpressing *draper*. Draper has three isoforms with distinct functions (Logan et al., 2012) – Draper-I regulates engulfment of axonal debris through its intracellular immunoreceptor tyrosine-based activation motif (ITAM); Draper-II is an inhibitor of Draper-I, reduces debris clearance when overexpressed, and is selectively expressed in adults; Draper-III lacks the ITAM and its function is unknown. Notably, Draper-I and -III are expressed in the larval brain and body wall (Logan et al., 2012). We overexpressed each isoform and examined cross-neuron plasticity. Here, we focused on MN6-Ib because in our previous study, this MN revealed a modest level of cross-neuron plasticity (Han et al., 2022; Wang et al., 2021), compared to the significantly higher plasticity at MN4-Ib which may not show further enhancement. To establish a baseline for MN6-Ib activity, we utilized GCaMP imaging together with electrophysiology recording to isolate the Ib EPSP (Figure S1C), and found that MN6-Ib contributes 53% of total EPSP.

Next, we assayed MN6-Ib structural and functional plasticity in Draper overexpressing animals. Elevating *draper-I* levels in either glial cells or muscles did not enhance Ib bouton growth upon Is ablation (Figure 4A and Figure S7A-F), but surprisingly, the EPSP amplitude and quantal content showed a significant increase (Figure 4B,C and Figure S7G-K). These results suggested that overexpressing *draper-I* can boost the Ib functional compensation. Unlike Draper-I, Draper-II is proposed to function as a repressor for debris engulfment. Indeed, we found that overexpressing *draper-II* in glial cells suppressed the compensation of Ib bouton number (Figure 4D and Figure S8A-D), as well as the compensation of EPSP and quantal content (Figure 4E and F, and Figure S8G-K). Interestingly, overexpressing *draper-II* in muscles suppressed the Ib structural compensation (Figure S8E and S8F), but not the functional compensation (Figure 4D-F). This data fit our previous observation that cross-neuron plasticity is primarily regulated by Draper activity in glial cells. Finally, overexpressing *draper-III* in either glial cells or muscles did not elicit Ib structural compensation (Figure 4G and Figure S9A-F) or EPSP (Figure 4H and Figure S9G-J), but muscle overexpression caused a slight increase of quantal content (Figure 4I and Figure S9K), suggesting that Draper-III can trigger cross-neuron plasticity when overexpressed post-synaptically. Taken together, our data suggested, 1) Draper-I is the functional isoform in glial cells for cross-neuron plasticity; 2) muscles are capable of triggering cross-neuron plasticity; 3) increasing *draper-I* or *-III* may be beneficial during neural degeneration.

**Figure 4.**
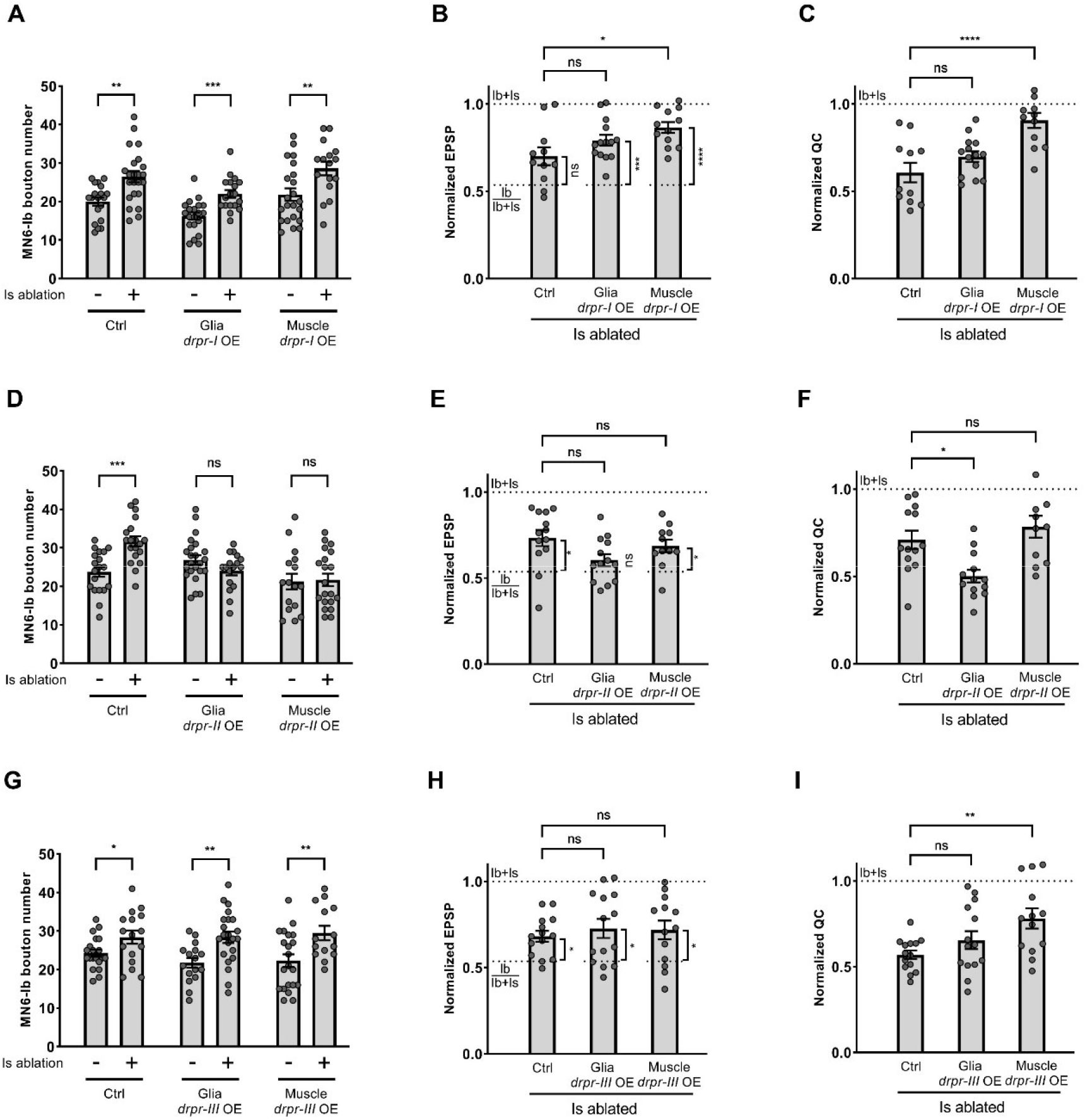
Overexpression of Draper-I boosts cross-neuron plasticity. A. Quantification of MN6-Ib bouton number in no ablation and Is ablated larvae in control, glia *draper-I* overexpression, and muscle *draper-I* overexpression backgrounds. Control (N = 19 and 23 NMJs), t(40)=3.493, p=0.0012, unpaired t-test. Glia *draper-I* overexpression (N = 20 and 18 NMJs), t(36)=4.103, p=0.0002, unpaired t-test. Muscle *draper-I* overexpression (N = 22 and 16 NMJs), t(36)=2.818, p=0.0078, unpaired t-test. B. Quantification of normalized EPSP of Is ablated larvae in control, glia *draper-I* overexpression, and muscle *draper-I* overexpression backgrounds. F(2, 34)=4.361, p=0.0206, one-way ANOVA. Is ablated control vs glia *draper-I* overexpression, p=0.2235. Is ablated control vs muscle *draper-I* overexpression, p=0.0153. Is ablated control vs Ib/Ib+Is, t(19)=2.074, p=0.0519, unpaired t-test. Is ablated in glia *draper-I* overexpression vs Ib/Ib+Is, t(22)=4.041, p=0.0005, unpaired t-test. Is ablated in muslce *draper-I* overexpression vs Ib/Ib+Is, t(20)=5.061, p<0.0001, unpaired t-test. C. Quantification of normalized quantal content of Is ablated larvae in control, glia *draper-I* overexpression, and muscle *draper-I* overexpression backgrounds. F(2, 34)=12.27, p<0.0001, one-way ANOVA. Is ablated control vs glia *draper-I* overexpression, p=0.2951. Is ablated control vs muscle *draper-I* overexpression, p<0.0001. For B and C, N (NMJs) = 11, 14, 12. D. Quantification of MN6-Ib bouton number in no ablation and Is ablated larvae in control, glia *draper-II* overexpression, and muscle *draper-II* overexpression backgrounds. Control (N = 19 and 19 NMJs), t(36)=4.319, p=0.0001, unpaired t-test. Glia *draper-II* overexpression (N = 23 and 17 NMJs), t(38)=1.626, p=0.1122, unpaired t- test. Muscle *draper-II* overexpression (N = 16 and 20 NMJs), t(34)=0.1763, p=0.8611, unpaired t-test. E. Quantification of normalized EPSP of Is ablated larvae in control, glia *draper-II* overexpression, and muscle *draper-II* overexpression backgrounds. F(2, 34)=2.731, p=0.0795, one-way ANOVA. Is ablated control vs glia *draper-II* overexpression, p=0.0680. Is ablated control vs muscle *draper-II* overexpression, p=0.7116. Is ablated control vs Ib/Ib+Is, t(21)=2.606, p=0.0165, unpaired t-test. Is ablated in glia *draper-II* overexpression vs Ib/Ib+Is, t(21)=1.008, p=0.3250, unpaired t-test. Is ablated in muscle *draper-II* overexpression vs Ib/Ib+Is, t(19)=2.162, p=0.0436, unpaired t-test. F. Quantification of normalized quantal content of Is ablated larvae in control, glia *draper-II* overexpression, and muscle *draper-II* overexpression backgrounds. F(2, 34)=8.539, p=0.0010, one-way ANOVA. Is ablated control vs glia *draper-II* overexpression, p=0.0130. Is ablated control vs muscle *draper-II* overexpression, p=0.5614. For E and F, N (NMJs) = 13, 13, 11. G. Quantification of MN6-Ib bouton number in no ablation and Is ablated larvae in control, glia *draper-III* overexpression, and muscle *draper-III* overexpression backgrounds. Control (N = 20 and 16 NMJs), t(23.64)=2.129, p=0.0439, unpaired t-test with Welch’s correction.. Glia *draper-III* overexpression (N = 16 and 22 NMJs), t(36)=3.261, p=0.0024, unpaired t-test. Muscle *draper-III* overexpression (N = 21 and 14 NMJs), t(33)=2.852, p=0.0074, unpaired t-test. H. Quantification of normalized EPSP of Is ablated larvae in control, glia *draper-III* overexpression, and muscle *draper-III* overexpression backgrounds. F(2, 38)=0.2634, p= 0.7698, one-way ANOVA. Is ablated control vs glia *draper-III* overexpression, p=0.7729. Is ablated control vs muscle *draper-III* overexpression, p=0.8508. Is ablated control vs Ib/Ib+Is, t(22)=2.291, p=0.0319, unpaired t-test. Is ablated in glia *draper-III* overexpression vs Ib/Ib+Is, t(22)=2.299, p=0.0314, unpaired t-test. Is ablated in muslce *draper-III* overexpression vs Ib/Ib+Is, t(21)=2.214, p=0.0380, unpaired t-test. I. Quantification of normalized quantal content of Is ablated larvae in control, glia *draper-III* overexpression, and muscle *draper-III* overexpression backgrounds. F(2, 38)=4.981, p=0.0120, one-way ANOVA. Is ablated control vs glia *draper-III* overexpression, p=0.4070. Is ablated control vs muscle *draper-III* overexpression, p=0.0089. For H and I, N (NMJs) = 14, 14, 13. Error bars indicate ± SEM, ns = non-significant, *p<0.05, **p<0.01, ***p<0.001, ****p<0.0001.

### Cross-neuron plasticity does not rely on Ib and Is co-innervation of muscles

In the experiments above, only muscles that are co-innervated by both Ib and Is MNs were analyzed (Figure S1A). Therefore, cell death and the axonal debris after Is MN ablation can be sensed by glial cells that wrap the axons as well as by the postsynaptic muscles. We reasoned that if glial cells are the major player to transmit the signal, then co-innervation of Ib and Is on the same muscle should not be required to induce cross-neuron plasticity upon Is ablation (i.e. glial contact is sufficient to induce cross-neuron plasticity). To test this hypothesis, we examined MN11-Ib which innervates muscle 11 without Is co-innervation. We found a significant increase of bouton number (Figure 5A- C), EPSP amplitude, and quantal content (Figure 5D-H) of MN11-Ib upon Is ablation, suggesting that even without a co-innervating Is MN on the muscle target, Ib MNs still respond to Is ablation and support our model that glial cells play an important role in this process. In addition, this result further demonstrated that cross-neuron plasticity is a general mechanism in multiple MNs.

**Figure 5.**
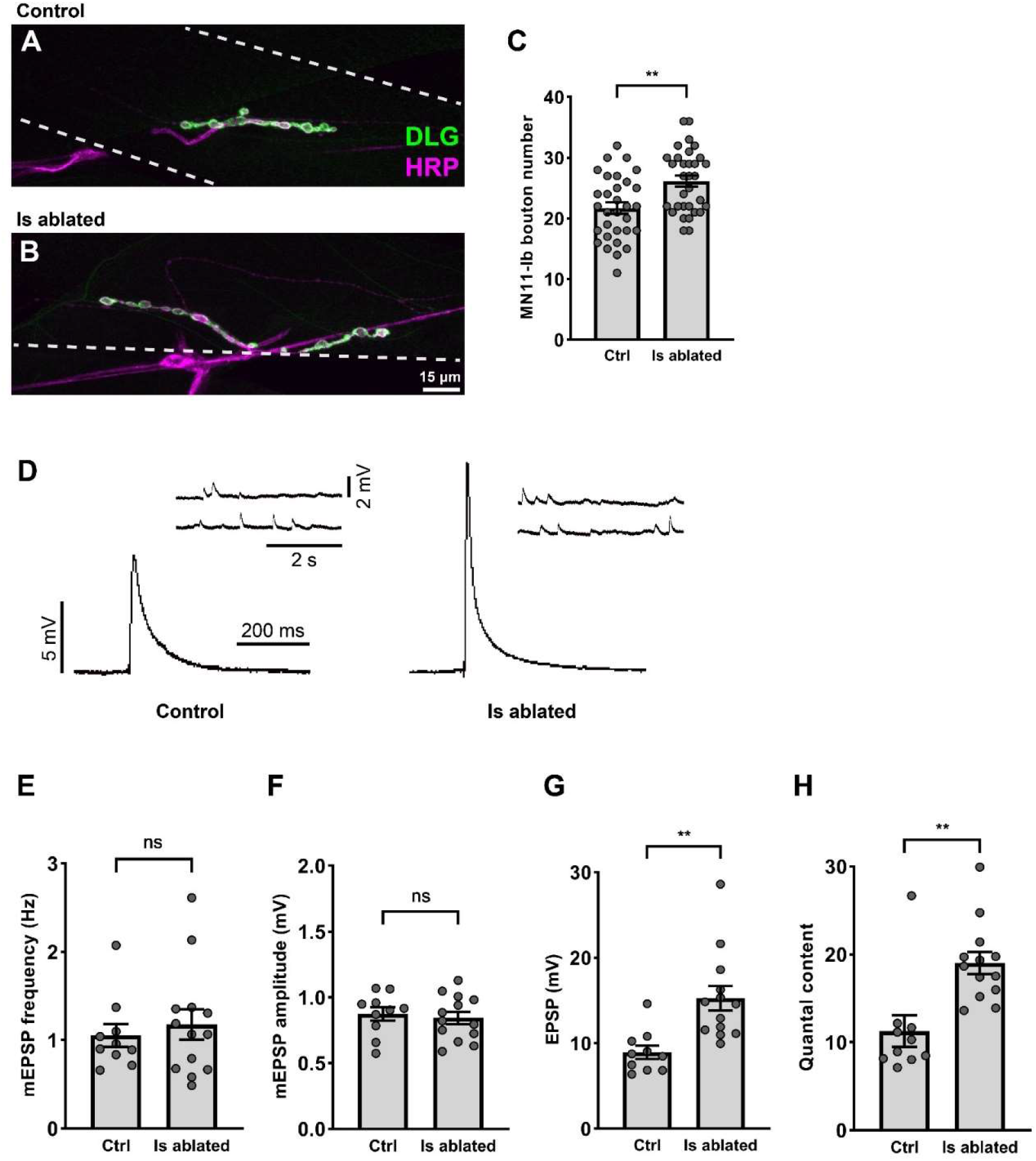
MN11-Ib showed cross-neuron plasticity upon Is ablation. A and B. NMJs of MN11-Ib in third instar control (*Is>GFP*) and Is ablated (*Is>GFP,hid,rpr*) larvae labeled with GFP (green) and HRP (magenta). Note the larger NMJs in Is ablated larvae. C. Quantification of MN11-Ib bouton number in no ablation (N = 31 NMJs) and Is ablated (N = 32 NMJs) larvae. t(61)=3.408, p=0.0012, unpaired t-test. D. EPSP and mEPSP recordings of muscle 11 from no ablation and Is ablated larvae. E. Quantification of mEPSP frequency. T(21)=0.5408, p=0.5943, unpaired t-test. F. Quantification of mEPSP amplitude. T(21)=0.4446, p=0.6611, unpaired t-test. G. Quantification of EPSP amplitude. T(18.02)=3.840, p=0.0012, unpaired t-test with Welch’s correction. H. Quantification of quantal content. T(21)=3.657, p=0.0015, unpaired t-test. For E-H, N (NMJs) = 10, 13. Error bars indicate ± SEM, ns = non-significant, **p<0.01.

### Acute Is MN ablation induces functional plasticity, but not structural plasticity

Nervous system plasticity generally declines as animals age (Ackerman et al., 2021; Purice et al., 2016). In our previous experiments, genetic ablation of Is MNs occurred in late embryonic stages when the animal is still developing, and synapses are undergoing extensive expansion and pruning (Wang et al., 2021). Therefore, it is important to understand whether cross-neuron plasticity is only inducible in early permissive stages, or if it is persists throughout development. To test if cross-neuron plasticity can be induced in later developmental stages, we established a heat-shock induced Is ablation system (Figure 6A). Animals were raised at 18°C and collected at different developmental stages from late stage embryos to third instar larvae and subjected to heat-shock. Heat-shock induces expression of a flippase transgene which removes a stop codon flanked by two FRT sites, thus allowing the *Is-GAL4* to drive the expression of cell death genes (Figure 6A). After Is ablation, animals were grown at 18°C and examined at late third instar.

**Figure 6.**
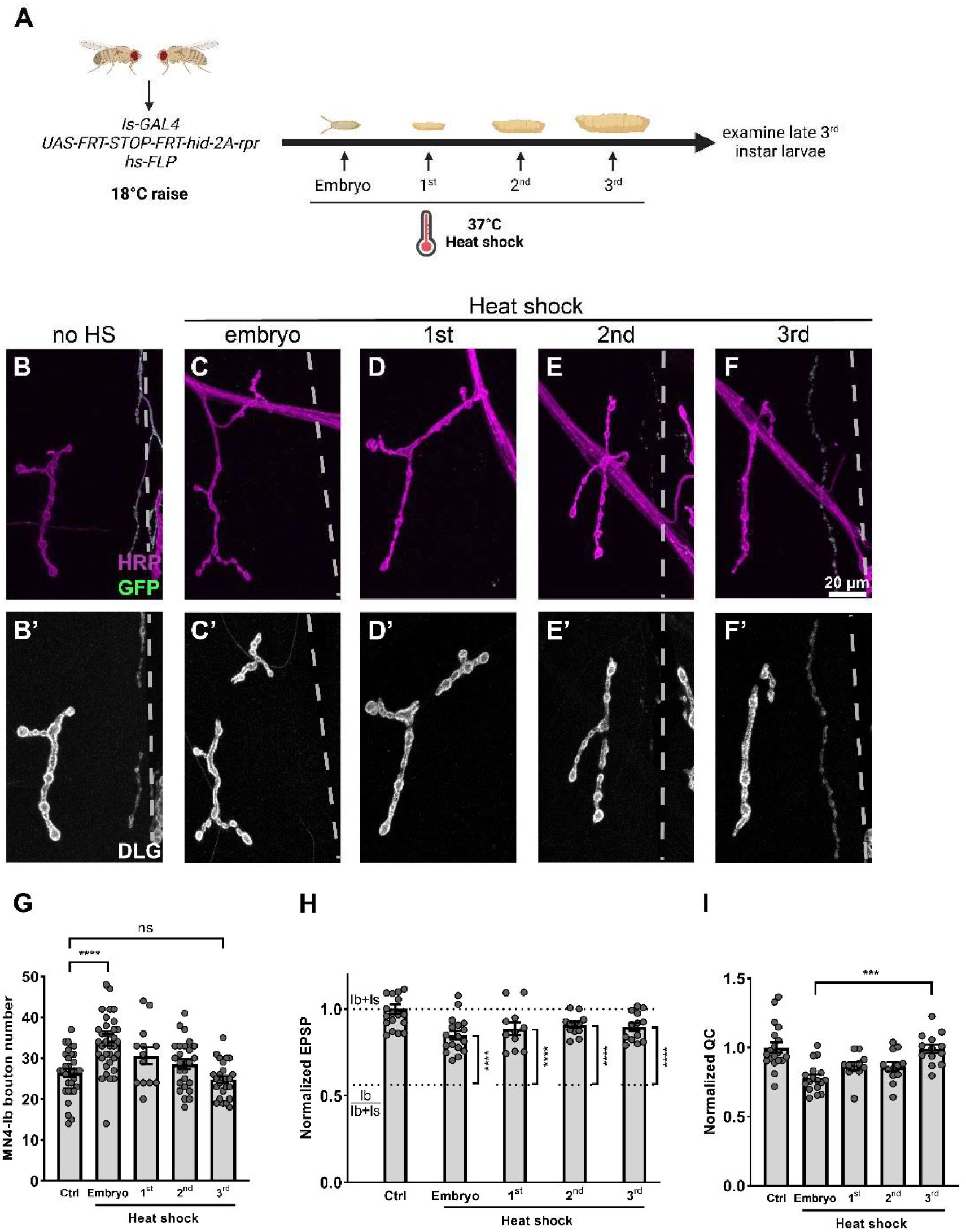
Acute Is MN ablation induces functional plasticity, but not structural plasticity. A. Schematic of heat-shock induced Is MN ablation. B-F and B’-F’. NMJs of MN4-Ib in late third instar larvae (*hs-FLP,UAS-GFP/+;UAS-FRT stop FRT-hid-2A-rpr/+;Is-GAL4/+*) with (B, B’) no heat-shock, (C, C’) embryo heat-shock, (D, D’) first instar heat-shock, (E, E’) second instar heat-shock and (F, F’) third instar heat-shock, stained with GFP (green), HRP (magenta), and DLG (grey). In embryos and first and second instar heat-shocked larvae, the Is NMJs were fully cleared, while Is synaptic debris remains in third instar heat-shocked larvae (F and F’). G. Quantification of MN4-Ib bouton number in late third instar larvae with Is MNs ablated at different developmental stages. F(4, 127)=10.23, p<0.0001, one-way ANOVA. Control vs embryo heat-shock, p<0.0001. Control vs first instar heat shock, p=0.3021. Control vs second instar heat-shock, p=0.7310. Control vs third instar heat-shock, p=0.8756. N (NMJs) = 33, 36, 13, 25, 25. H. Comparison of the normalized EPSP of late third instar larvae with Is MNs ablated at different developmental stages to Ib/Ib+Is baseline. Embryo heat-shock, t(27)=7.126, p<0.0001, unpaired t-test. First instar heat-shock, t(21)=6.622, p<0.0001, unpaired t-test. Second instar heat-shock, t(22)=9.485, p<0.0001, unpaired t-test. Third instar heat-shock, t(23)=8.557, p<0.0001, unpaired t-test. I. Quantification of normalized quantal content of late third instar larvae with Is ablated at different developmental stages. F(4, 67)=9.109, p<0.0001, one-way ANOVA. Embryo heat-shock vs third instar heat-shock, p=0.0002. Non-significant for the others. This result suggested an increase of cross-neuron plasticity when acutely ablated Is MNs. For H and I, N (NMJs) = 19, 17, 11, 12, 13. Error bars indicate ± SEM, ns = non-significant, ***p<0.001, ****p<0.0001.

We first calculated the ablation efficiency and confirmed that our approach ablated approximately 80% of Is MNs, providing sufficient samples for analysis (Figure S10). We then examined the NMJs and VNCs of animals with Is MNs ablated at different time points and confirmed that all debris from ablated Is MNs was removed by the time we assayed, which allows us to faithfully study the Ib responses (Figure 6B-F and Figure S11A-E). Examining muscle 4s, we found a significant MN4-Ib bouton number increase only when Is MNs were ablated in late embryonic stages (Figure 6B, C and G), but not in animals with later ablation (Figure 6D-G). However, despite a lack of structural changes, MN4-Ib EPSP and quantal content were increased in all stages upon Is ablation (Figure 6H and 6I). These results suggest that 1) acute Is MN ablation induces functional plasticity of Ib MNs; 2) structural and functional plasticity may be regulated by different mechanisms; and 3) functional plasticity is not simply a consequence of more boutons.

### Cross-neuron plasticity enhances larval locomotion

In *Drosophila*, the Ib MNs are considered tonic neurons that provide sustained responses, while the Is MNs are phasic neurons that respond and adapt quickly (Aponte-Santiago and Littleton, 2020). It has been thought that the tonic Ib MNs are the major drive for normal larval behavior such as forging and crawling, whereas the phasic Is MNs are responsible for quick actions such as escaping (Schaefer et al., 2010). Here, we wondered what the behavioral consequences would be upon Is ablation, focusing on behaviors that are attributed to both Ib and Is activity. First, we analyzed the crawling behavior in freely moving larvae. Interestingly, we found that Is ablated animals had a faster crawling speed but turned less compared to wild type controls (Figure 7 A-C). We reasoned that the faster crawling speed is a consequence of Ib elevated activity upon Is ablation whereas the turning defect is due to the loss of multiple-targeted phasic Is MNs to aid synchronous movement to one side. Next, we analyzed larval rolling behavior, a well-characterized escape behavior. When wild type larvae are immersed in water on heat plate, a stereotyped rolling behavioral response is readily triggered (Chattopadhyay et al., 2012) (Figure 7D, E). We found that wild type larvae displayed sustained roll behavior, whereas Is ablated larvae showed extensive head stretch and bends but failed to perform complete rolling (Figure 7E and Video 1). This result suggested that the Is MNs play an important role in the escape neural circuit, and cross-neuron plasticity cannot compensate the behavioral changes, as expected to their distinct circuit partners.

**Figure 7.**
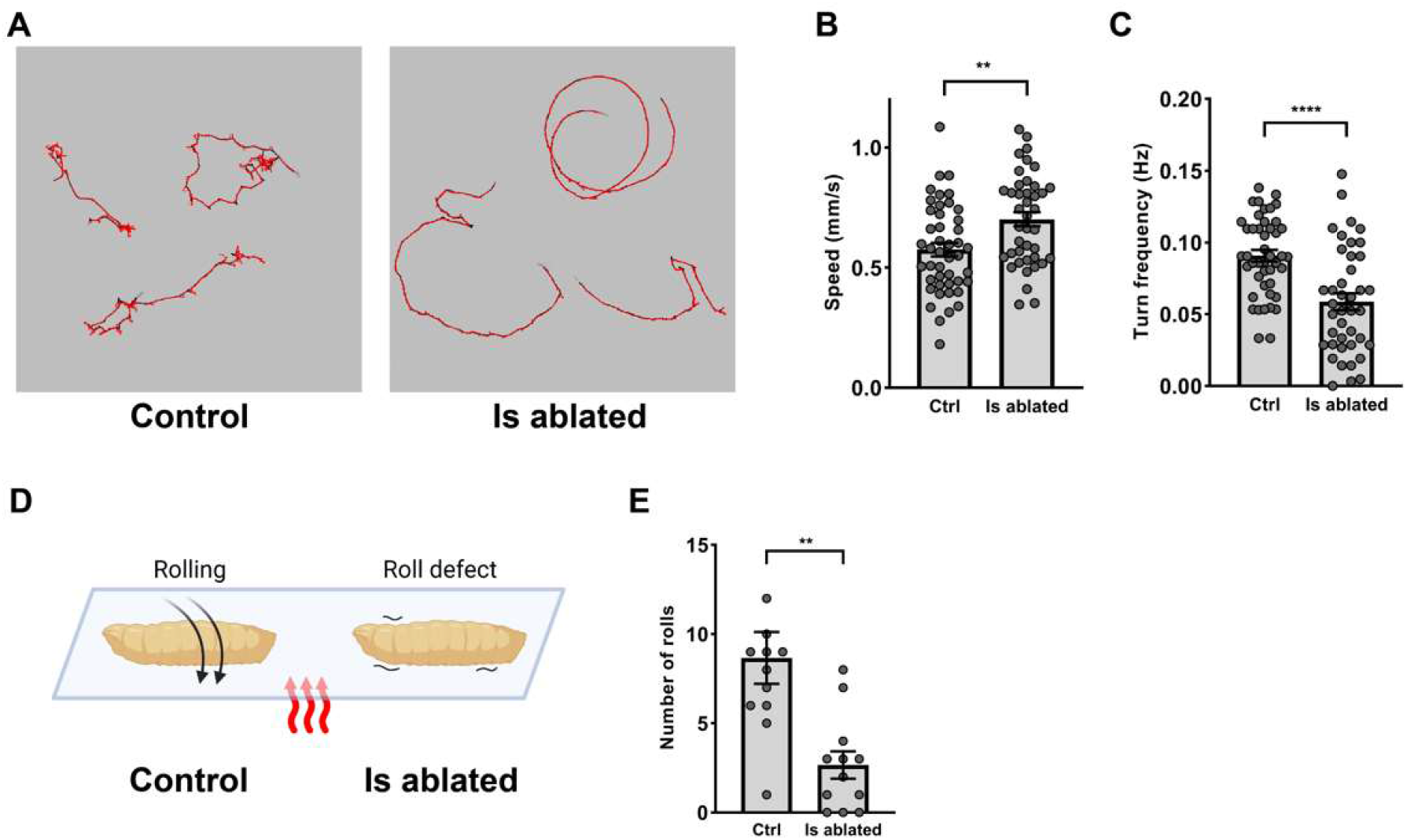
Cross-neuron plasticity bolsters larval locomotion speed. A. Representative crawling traces of control and Is ablated larvae. Note the control larvae perform more turns than Is ablated larvae. B. Quantification of crawling speed of control and Is ablated larvae. t(84)=3.107, p=0.0026. C. Quantification of turn frequency of control and Is ablated larvae. t(72.67)=4.518, p<0.0001. For C and D, N (larvae) = 45, 41. D. Schematic of heat induced roll behavior difference in control and Is ablated larvae. E. Quantification of the number of rolls of control and Is ablated larvae. t(16.59)=3.647, p=0.0021. N (larvae) = 12, 12. Error bars indicate ± SEM, **p<0.01, ****p<0.0001.

## Discussion

Neuronal cell death is a hallmark of aging, injury, and many neurodegenerative diseases and significant efforts have been made to delay, prevent, or ameliorate these incidents. Most neurons cannot regenerate to restore function, but healthy bystander neurons may provide an alternative to overcome dysfunction after neuronal cell death. Indeed, several studies have reported that bystander neurons enhance their structurally and functionally properties when they detect loss of neighboring neurons (Aponte-Santiago et al., 2020; Wang et al., 2021). This attractive model highlights a new type of plasticity of the nervous system to maintain its homeostatic state, which we termed as cross-neuron plasticity.

In this study, we found that the *Drosophila* engulfment receptor, Draper, and an interacting kinase, Shark, are required in glial cells for cross-neuron plasticity in Ib MNs after loss of Is MNs. Overexpression of *draper* boosted the compensatory changes in bystander Ib MNs, providing an exciting avenue to recover the functional defects. We also examined induction of cross-neuron plasticity at different time points and found that functional plasticity is inducible at all larval stages, suggesting cross-neuron plasticity does not simply reflect a highly plastic temporal window during embryonic development. Taken together, our study revealed important mechanistic insights into cross-neuron plasticity and establish an entry point to understand how healthy bystander neurons respond to their dying neighbors.

### Role of Draper/Shark pathway in glia-neuron crosstalk

Neuronal debris generated during development or injury must be removed in a timely manner to ensure it does not damage other tissues. Degenerating neurons express or secrete “eat me” signals that are recognized by engulfment receptors on phagocytic cells, including glial cells (Ji et al., 2022; McLaughlin et al., 2019). In the *Drosophila* nervous system, the engulfment receptor, Draper, interacts with several “eat me” signals, including SIMU (Kurant et al., 2008), pretaporter (Kuraishi et al., 2009), and phosphatidylserine (Sapar et al., 2018; Tung et al., 2013). Upon ligand binding, Draper is phosphorylated by Src42a and binds to Shark, which together become the signaling core for the clearance pathway (Ziegenfuss et al., 2008). Draper/Shark first activate Rac1 through DRK/DOS/SOS or dCed-12/MBC/Crk complexes to initiate glial membrane recruitment to engulf the debris from degenerating neurons (Lu et al., 2014; Ziegenfuss et al., 2012). In parallel, Draper/Shark activate the dJNK pathway and downstream dAP-1 and STAT92E transcription factors to drive the expression of engulfment genes (Doherty et al., 2014; Lu et al., 2017). This phagocytic pathway has been extensively studied during synaptic pruning and injury induced axon degeneration, and here we implicate Draper/Shark in removing neuronal debris during programmed cell death. As transcription factors of the vertebrate innate immune system, AP-1 and STATs regulate many essential cellular processes such as differentiation, proliferation, apoptosis, and expression of inflammatory cytokines and chemokines, which regulate the behavior and function of other cells in response to pathogens or damages (Hess et al., 2004; Levy and Darnell, 2002). Utilizing severed sensory neuron axons in adult fly wings, a recent study revealed that glial cells function through Draper->JNK->dAP-1 to suppress axon transport in bystander neurons, suggesting that glial cells detect and spread an injury signal to nearby healthy neurons (Hsu et al., 2020). Here, we explored this crosstalk in the neuromuscular system and found that cross-neuron plasticity in bystander neurons also relies on Draper signaling. Loss of either *draper* or *shark* suppressed cross-neuron plasticity, and *draper* overexpression boosted the plasticity. Our results provide a link between the injury-induced glial response and the plasticity changes in bystander neurons and further suggest a homeostatic role of the Draper/Shark pathway when neuronal cell death happens. These positive effects seemingly contradict the Draper/Shark-mediated axon transport defects in bystander neurons when severing sensory neurons in the adult fly wing (Hsu et al., 2020). The opposing effects might be due to different mechanisms acting in sensory and motor systems, or more likely, to the different time length after ablation/injury. In the wing, axon transport defects were observed 3 hours after injury, whereas we examined cross-neuron plasticity at least two days after ablation. Indeed, the behavioral defects caused by wing injury fully recovered after 6 hours, suggesting the bystanders may eventually compensate for injured neurons. Nevertheless, our data support a model whereby debris from dead/injured neurons triggers the Draper/Shark pathway in surrounding glial cells to transmit a signal to healthy bystander neurons to compensate the functional loss.

### Distinct mechanisms may instruct structural and functional plasticity

In this study, we showed that the Draper/Shark pathway is required for cross-neuron plasticity, but important questions remain – what is the signal from glial cells to bystander neurons, and how do bystander neurons respond to such a signal to alter their morphology and functional output? Glial cells are intimately associated with neurons offering several opportunities for communication. First, glial cells and neurons may directly interact through cell surface receptors/ligands, such as Notch/Delta (Calderon et al., 2022) and Nrx/Nlg (Ackerman et al., 2021). Alternatively, glial cells may secrete molecules, like Wnt (Simões et al., 2022), Spz (McLaughlin et al., 2019), and others, that bind to corresponding receptors on bystander neurons. It will be of great interest to examine if any of these signaling pathways are involved in cross-neuron plasticity. Notably, several lines of evidence suggest that bystander neurons may utilize distinct mechanisms for the structural and functional components of cross-neuron plasticity. In our previous study, MN12-Ib displayed robust structural plasticity but no functional plasticity when the Is neighbor was ablated, unlike MN6-Ib and MN4-Ib which showed structural and functional plasticity (Wang et al., 2021). In the current study, when we acutely ablated Is MNs in third instar larvae, we only observed an increase of the EPSP, but no correlated structural bouton increase. Together these data suggest several non-exclusive models: 1) downstream of Draper/Shark, different mechanisms may be utilized to regulate structural or functional cross-neuron plasticity, 2) different MNs have differing capabilities to show either, or both plasticity changes, and 3) structural plasticity may require time to add boutons gradually, whereas the functional plasticity may utilize established synaptic architecture to achieve a fast response. Indeed, active zones are highly dynamic and can acutely alter synaptic strength without the need of additional boutons. For example, the synaptic machinery, such as presynaptic active zone structural proteins (Goel et al., 2017; Gratz et al., 2019; Weyhersmüller et al., 2011) or postsynaptic neurotransmitter receptors (Schmid et al., 2008), could increase their density to elevate synaptic release or the response to each synaptic vesicle, respectively. In addition, the properties of individual active zones, such as the size of the readily releasable pool and the release probability, may increase following perturbations (Kiragasi et al., 2017; Li et al., 2018; Müller et al., 2015). Notably, in our previous study, the synaptic machineries in bystander neurons were not significantly altered, suggesting that functional cross-neuron plasticity may modify synaptic properties (Wang et al., 2021). Examining these synaptic parameters will provide hints to explain the functional changes in bystander neurons.

### Cross-neuron plasticity provides a novel mechanism enhance circuit activity

Neuronal cell death is a hallmark in aging, injury, and neurodegenerative diseases, but the nervous system possesses mechanisms to compensate for the loss of circuit members. In this study, we examined cross-neuron plasticity, a mechanism where healthy bystander neurons detect and respond to death of neighboring neurons. Substantial evidence supports the potency of cross-neuron plasticity. For example, denervating muscles in both vertebrates and invertebrates lead to compensatory axon terminal expansion from healthy bystander neurons, which may eventually provide functional recovery (Aponte-Santiago et al., 2020; Brown et al., 1981; Edds, 1953; Han et al., 2022; Wang et al., 2021). On the behavioral level, severing sensory neurons in the adult fly wing causes immediate sensory defects that are rescued within 6 hours (Hsu et al., 2020). In humans, sensory loss can lead to adaptation of brain circuits to utilize the remaining senses to navigate, a process known as cross-modal recruitment and compensatory plasticity (or cross-modal plasticity) (Lazzouni and Lepore, 2014; Lee and Whitt, 2015). Although the mechanisms underlying these phenotypes may vary, together they represent an intriguing, alternative paradigm that the nervous system can utilize to counteract synaptic dysfunction and neuronal death.

## Methods

### Fly and antibody reagents

Fly resources

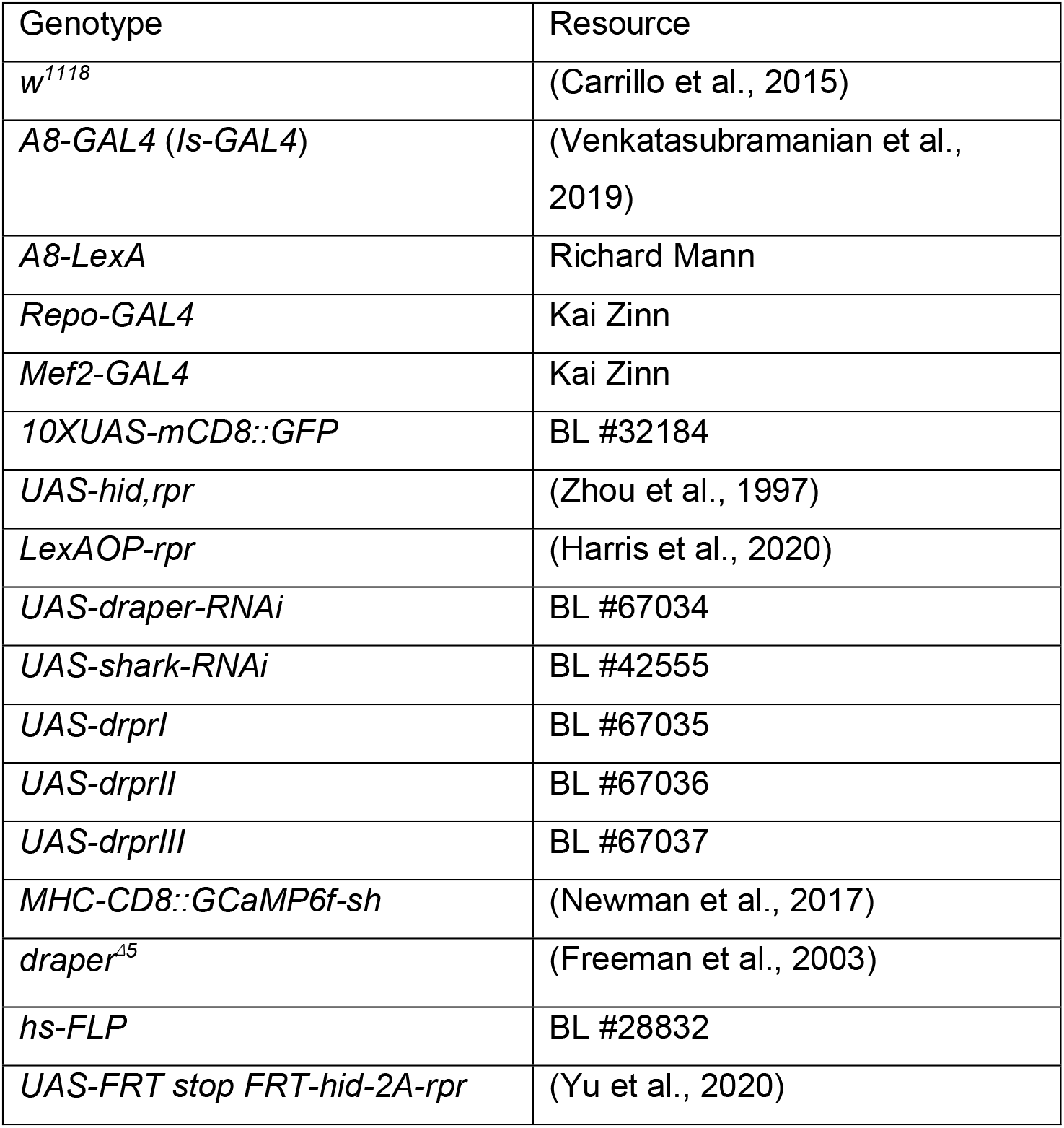

### Antibody resources

| Antibody | Source | Concentration |
| --- | --- | --- |
| Mouse anti-Repo | Developmental Studies Hybridoma Bank, #8D12 | 1:50 |
| Chicken anti-GFP | Michael Glotzer Lab | 1:10,000 |
| Rabbit anti-GFP | Thermo Fisher Scientific, #A11122 | 1:500 |
| Mouse anti-DLG | Developmental Studies Hybridoma Bank, #4F3 | 1:100 |
| Rabbit anti-DLG | (Koh et al., 1999) | 1:40,000 |
| Mouse anti-Draper | Developmental Studies Hybridoma Bank, #8A1-S | 1:30 |
| Goat anti-HRP-Alexa Fluor 405 | Jackson ImmunoResearch, #123-475-021 | 1:100 |
| Goat anti-HRP-TRITC | Jackson ImmunoResearch, #123-025-021 | 1:100 |
| Goat anti-HRP-Alexa Fluor 647 | Jackson ImmunoResearch, #123-605-021 | 1:100 |
| Goat anti-mouse-Alexa Fluor 647 | Thermo Fisher Scientific, #A32728 | 1:500 |
| Goat anti-chicken-Alexa Fluor 488 | Thermo Fisher Scientific, #A11039 | 1:500 |
| Goat anti-rabbit-Alexa Fluor 568 | Thermo Fisher Scientific, #A11036 | 1:500 |
| Goat anti-rabbit-Alexa Fluor 488 | Thermo Fisher Scientific, #A11008 | 1:500 |

### Fly husbandry

*Drosophila* stocks and crosses were maintained at 25 °C except for the fly lines required for the heat-shock experiments. For crosses other than the heat-shock experiments, 6-8 females were mated with 3-5 males and flipped every day to ensure proper larval density. For heat-shock experiments, flies were mated and kept at 18 °C, and embryos or larvae at different developmental stages were collected for a 5 min 37 °C heat-shock in 1.5 mL Eppendorf tubes. Heat-shocked animals were transferred back to 18 °C until larvae were collected for analyses. Parental animals and larvae were randomly selected for mating and recording, respectively. We included *Is>GFP* in all experiments to ensure that the muscle 4s without Is ablation were co-innervated by both Ib and Is MNs, as approximately 20% of muscle 4s lack Is innervation naturally (Ashley et al., 2019). Both genders were equally used in this study.

### Dissections, immunocytochemistry, and imaging

Dissections and immunostaining were performed as previously described (Lobb-Rabe et al., 2022). Briefly, wandering third instar larvae were collected and dissected on sylgard plate in PBS. Samples were fixed by 4% paraformaldehyde for 20 min and then washed three times in PBT (PBS with 0.05% Triton X-100) for 15 min each. Samples were then blocked for 1 h in 5% goat serum (5% goat serum diluted in PBT) and incubated with primary antibodies at 4 °C overnight. Next, primary antibodies were washed out and secondary antibodies were applied at room temperature for 2 h. Finally, samples were washed and mounted in Vectashield (Vector Laboratories). Images were acquired on a Zeiss LSM800 confocal microscope using either a 40X plan-neofluar 1.3 NA objective, or a 63X plan-apo 1.4 NA objective. The same imaging parameters were applied to samples from the same set of experiments. Images were then analyzed and processed in ImageJ.

### Electrophysiology

Third instar larvae were dissected in magnetic chambers using modified HL3 saline (70 mM NaCl, 5 mM KCl, 10 mM MgCl2, 10 mM NaHCO3, 5 mM trehelose, 115 mM sucrose, 5 mM HEPES) with 0.5 mM calcium (Meng et al., 2020). Segmental nerves were severed near the VNC and the brain and VNC were removed to prevent endogenous action potentials. Samples were examined under a Nikon FS microscope with a 40X long-working distance objective to locate the muscle fibers and axons. Muscles 4, 6 or 11 from abdominal segment A3 and A4 were chosen and impaled by a 10–30 MΩ sharp electrode filled with 3 M KCl. To elicit EPSPs from MN6-Ib, the entire segmental nerve was drawn into a suction electrode, whereas for MN4-Ib and MN11-Ib, the intersegmental nerve above muscle 5 was stimulated. Each sample was first recorded for 1 min for miniature EPSPs (mEPSPs), followed by 2 min recordings of EPSPs at 0.2 Hz. 24 EPSPs were elicited and the larger 12 EPSPs were averaged to represent the mean EPSP. Due to the nonlinear summation of quantal content of large EPSPs, we corrected EPSP amplitude by the equations defined by (Martin, 1955) and elaborated by (Feeney et al., 1998). Quantal content (QC) was calculated by dividing the corrected EPSP amplitude by the mEPSP amplitude. Electrophysiology signals was amplified by a MultiClamp 700B amplifier (Molecular Devices), digitized with a Digidata 1550B (Molecular Devices), and acquired in pCLAMP 10 software (Molecular Devices). Axon stimulation was delivered by a Master-9 stimulator (A.M.P.I.). Data was finally analyzed with Mini Analysis software (Synaptosoft).

### GCaMP imaging coupled with electrophysiology

Third instar *MHC-CD8::GCaMP6f-Sh* larvae were dissected and processed as described above. Larval fillets were visualized under a 40X microscope and the GCaMP-positive Ib and Is NMJs were illuminated with an Aura II solid-state illuminator. Together with electrophysiology recordings, real time NMJ firing movies were recorded using a PCO Edge 4.2 camera and NIS-Elements Imaging Software. Due to the different evoked thresholds of Ib and Is MNs, stimulating voltages were fine tuned to isolate Ib MN firing and Ib+Is firing. EPSPs corresponding to specific MN firing combainations were categorized as Ib EPSP and Ib+Is EPSP. For each sample, Ib EPSP and Ib+Is EPSP were both recorded and the contribution of Ib MNs in wild type animals was calculated (Ib EPSP/Ib+Is EPSP).

### Behavioral assay

For the crawling assay, a third instar larva was placed on 2% agarose gel in a 100 mm petri dish at room temperature. Crawling trajectory was captured with a PiVR setup (Tadres and Louis, 2020) at 50 FPS and the centroids of the larva were considered as larval positions. Sampling frames were selected every 50 frames and the total travel distance was the sum of the distance between every two larval positions in consecutive samples frames. Total travel distance was divided by time to calculate speed. A larva was considered to turn if the angle between two vectors formed by connecting three larval positions in consecutive sampling frames is between 45 degrees and 150 degrees, and turn frequency was calculated by dividing the number of turns by time. For rolling behavior, we followed the “Global Heat Plate Assay” protocol (Chattopadhyay et al., 2012). Briefly, a third larva was placed in 200 μl water at the center of a plastic 35 mm petri dish at room temperature and then transferred onto a 95 °C heat block to elicit rolling behavior. Rolling behavior movies were captured by Point Grey FlyCap2 software. Only full lateral 360 degree rolls were counted by observing the dorsal trachea disappear under the larvae and emerge again on the opposite side.

### Statistics and reproducibility

All statistical analyses were performed using Prism 8 and mean and Standard Error of the Mean (SEM) were reported. Experimenters were blinded when performing bouton counts, and bouton numbers were linked with genotypes after counting. For other recordings, data was recorded by a computer and no subjective rating of data was involved. Measurements were recorded from distinct samples. Sample sizes were determined by existing studies in the field to enable statistical analyses and reproducibility. In general, for each experiment, at least 10 total data points were collected from each genotype. All experiments were performed with at least two independent biological replicates and data points were pulled and analyzed together. No data were excluded. All data were assumed to follow a Gaussian distribution. When comparing the no ablation versus Is ablated larvae under the same genetic background, two-sided Student’s t-test was used (Welch’s correction was used in case of unequal variance). When comparing across multiple conditions, one-way ANOVA with Turkey’s test was performed.

### Data Availability

The data that support the finding of this study are available from YW or RAC upon reasonable request. The behavioral data and the customized python code for data analyzation can be found at https://github.com/sihaohuanguc/larva_trajectory_process.

## Supporting information

Video 1. Rolling behavior

## Acknowledgement

This work is supported by NSF IOS-2048080, NINDS R01 NS123439 01, and a UChicago Faculty Diversity Grant to R.A.C, F31NS120458 and T32 GM007183 to M.L.R, and NIH R01NS070644 to Richard S. Mann (for support of L.V.). This work is also supported by funds from UChicago Biological Science Division, Committee of Developmental Biology and Department of Molecular Genetics & Cellular Biology. We thank the Bloomington Drosophila Stock Center (NIH P40OD018537) for fly lines. The monoclonal antibodies 4F3 and 8B12 were developed by Goodman, C., and the 8A1-S antibody was developed by Mary Logan, and they were obtained from the Developmental Studies Hybridoma Bank, created by the NICHD of the NIH and maintained at the University of Iowa, Department of Biology. We would like to thank Richard Mann (Columbia University), Robin E. Harris (Arizona State University) and Jean-Paul Vincent (the Francis Crick Institute) for sharing fly lines. We would also like to thank Richard Fehon, Ellie Heckscher, David Pincus, Audrina Daisy, Viola Nawrocka, Parisa Tajalli Tehrani Valverde and members from the Carrillo laboratory for valuable discussions and comments.

## Author contribution

Y.W. and R.A.C designed research; Y.W., R. Z., P.T.T.V, L.V., and M.L.R. performed experiments; Y.W., S. H. and J.A. analyzed data; Y.W. wrote the manuscript and J.A., M.L.R. and R.A.C. edited the manuscript.

---

**Figure S1.**
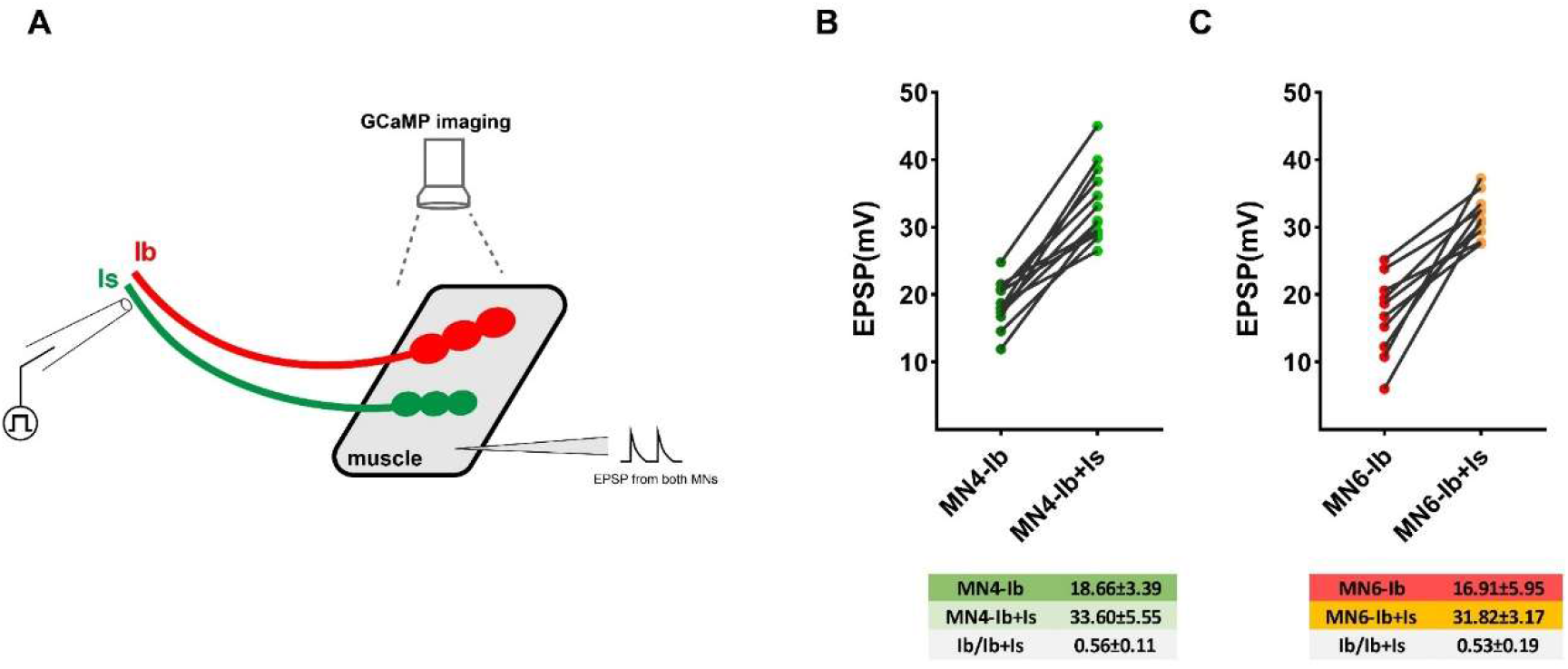
GCaMP imaging combined with electrophysiology recording to separate Ib and Is MN activity. A. Schematic of experimental setup. A GCaMP movie was recorded at the same time as recording EPSPs. B. Paired MN4-Ib and MN4-Ib+Is EPSP amplitudes. C. Paired MN6-Ib and MN6-Ib+Is EPSP amplitudes.

**Figure S2.**
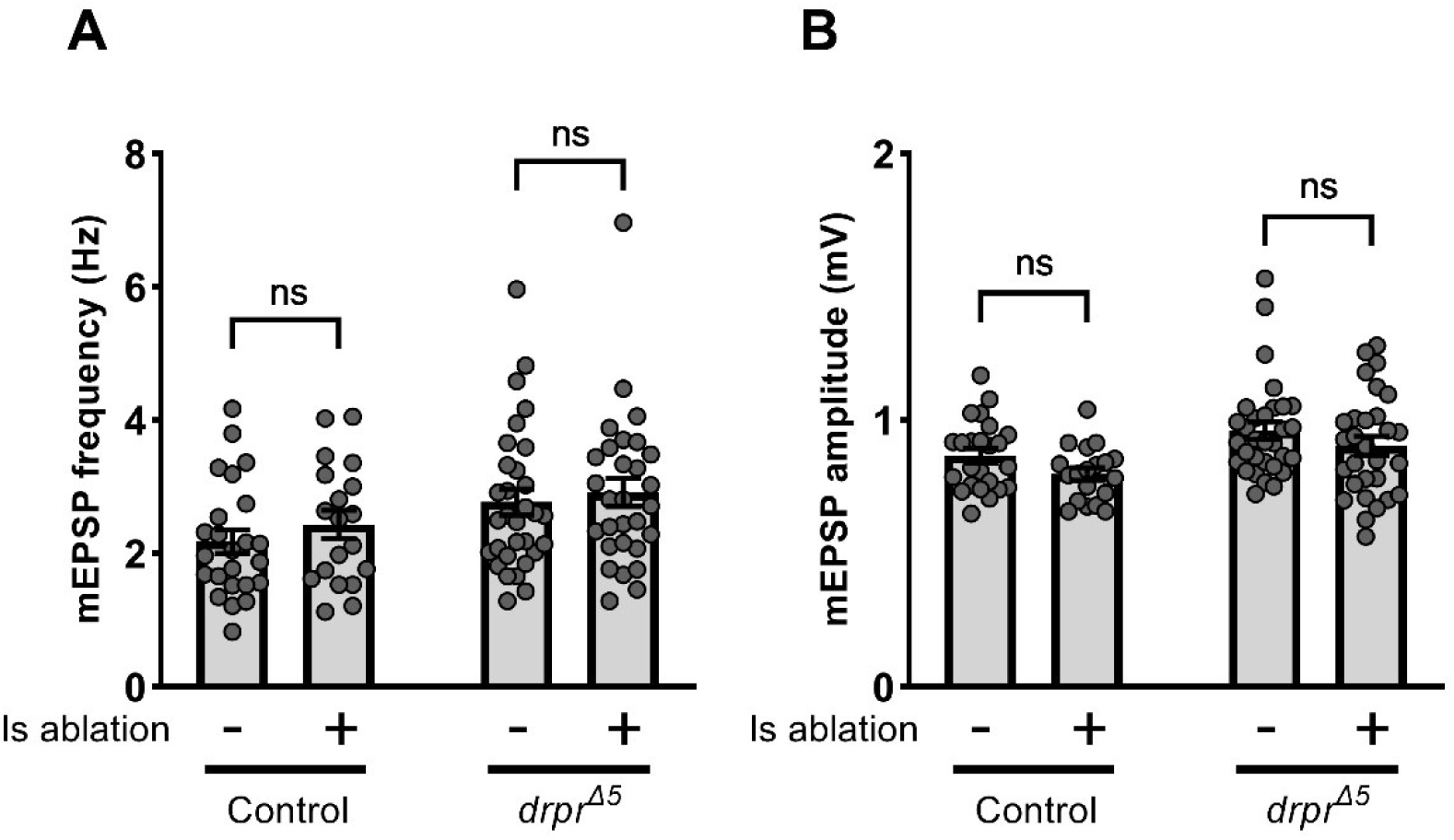
Quantification of mEPSPs of no ablation and Is ablated larvae in control and *drpr^Δ5^* backgrounds. A. Quantification of mEPSP frequencies in no ablation and Is ablated larvae in control and *drpr^Δ5^* backgrounds. Control, t(41)=0.9502, p=0.3476, unpaired t-test. *drpr^Δ5^*, t(58)=0.5102, p=0.6118, unpaired t-test. B. Quantification of mEPSP amplitude in no ablation and Is ablated larvae in control and *drpr^Δ5^* backgrounds. Control, t(41)=1.880, p=0.0672, unpaired t-test. *drpr^Δ5^*, t(58)=1.169, p= 0.2471, unpaired t-test. For A and B, N (NMJs) = 24, 19, 31, 29. Error bars indicate ± SEM, ns = non-significant.

**Figure S3.**
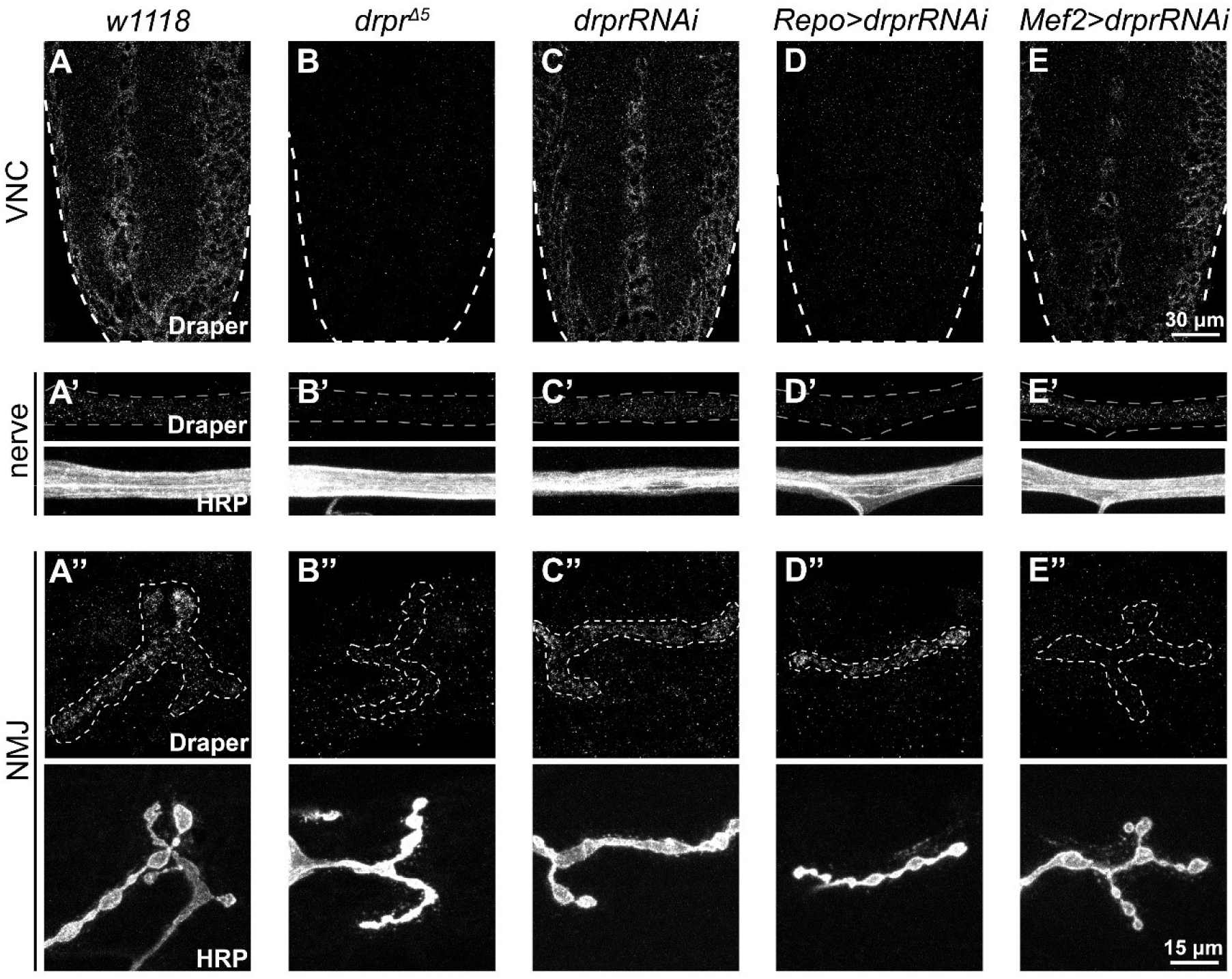
Validating draper RNAi efficiency. VNC, segmental nerves, and NMJs of (A, A’, A’’) *w1118*, (B, B’, B’’) *draper* mutant (*drpr^Δ5^*), (C, C’, C’’) *drprRNAi*, (D, D’, D’’) glia *draper* knockdown (*Repo>drprRNAi*), and (E, E’, E’’) muscle *draper* knockdown (*Mef2>drprRNAi*), stained with Draper and HRP. Note that glia *draper* knockdown completely removed the Draper signal from the VNC and segmental nerve, and muscle *draper* knockdown completely removed the Draper signal at the NMJs.

**Figure S4.**
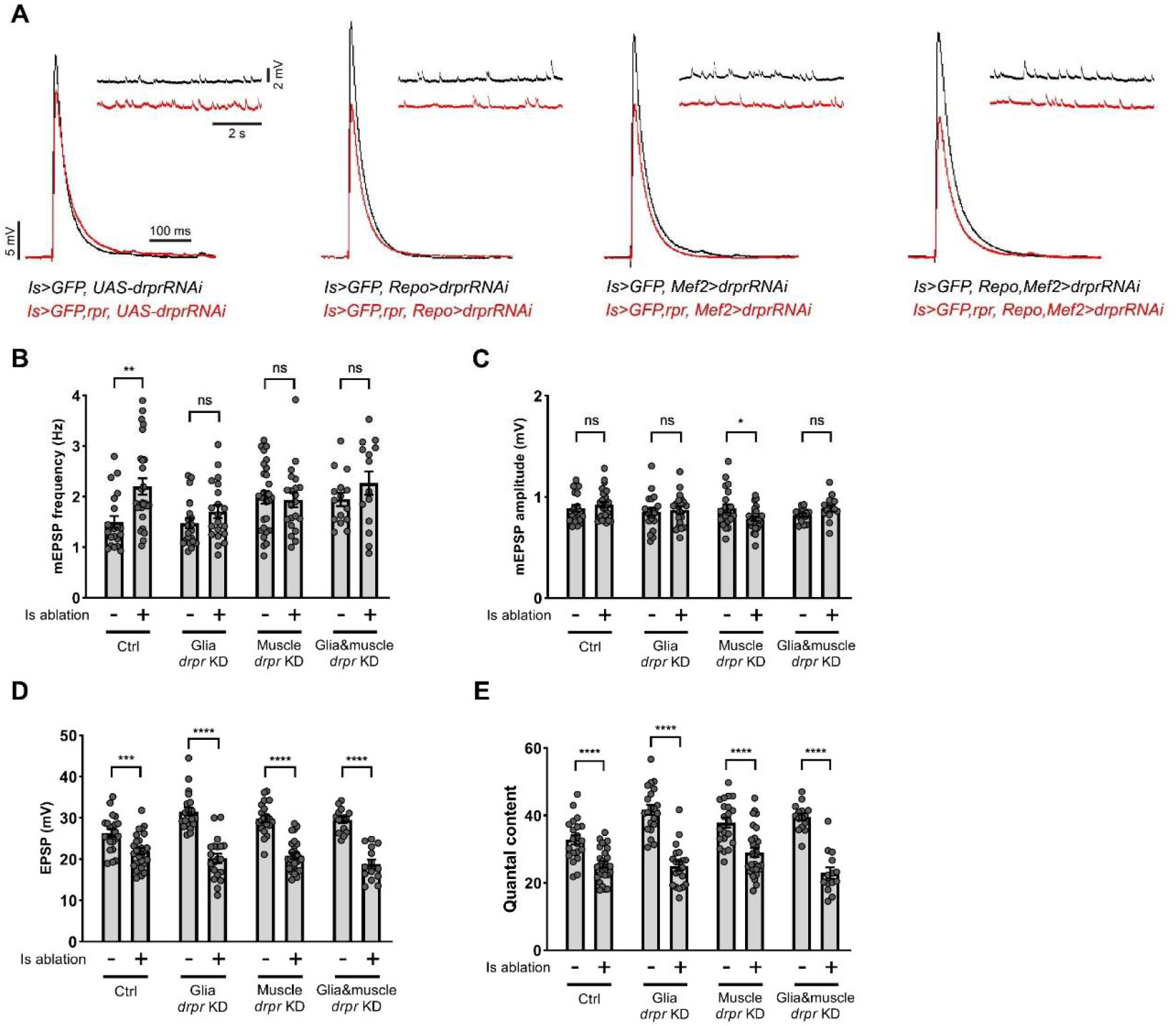
Electrophysiology recordings in cell type specific *draper* knockdown larvae. A. EPSP and mEPSP recordings from no ablation and Is ablated larvae in control, glia *draper* knockdown, muscle *draper* knockdown, and double knockdown backgrounds. B. Quantification of mEPSP frequency of no ablation and Is ablated larvae in control, glia *draper* knockdown, muscle *draper* knockdown, and double knockdown backgrounds. Control, t(45)=3.352, p=0.0016, unpaired t-test. Glia *draper* knockdown, t(39)=1.429, p=0.1610, unpaired t-test. Muscle *draper* knockdown, t(47)=0.2739, p=0.7854, unpaired t-test. Double knockdown, t(27)=1.219, p=0.2333, unpaired t-test. C. Quantification of mEPSP amplitude of no ablation and Is ablated larvae in control, glia *draper* knockdown, muscle *draper* knockdown, and double knockdown backgrounds. Control, t(45)=0.8366, p=0.4072, unpaired t-test. Glia *draper* knockdown, t(39)=0.3657, p=0.7165, unpaired t-test. Muscle *draper* knockdown, t(29.93)=2.279, p=0.0300, unpaired t-test with Welch’s correction. Double knockdown, t(20.18)=1.848, p=0.0793, unpaired t-test Welch’s correction. D. Quantification of EPSP amplitude of no ablation and Is ablated larvae in control, glia *draper* knockdown, muscle *draper* knockdown, and double knockdown backgrounds. Control, t(45)=3.579, p=0.0008, unpaired t-test. Glia *draper* knockdown, t(39)=7.543, p<0.0001, unpaired t-test. Muscle *draper* knockdown, t(47)=8.289, p<0.0001, unpaired t- test. Double knockdown, t(27)=8.247, p<0.0001, unpaired t-test. E. Quantification of quantal content of no ablation and Is ablated larvae in control, glia *draper* knockdown, muscle *draper* knockdown, and double knockdown backgrounds. Control, t(45)=4.472, p<0.0001, unpaired t-test. Glia *draper* knockdown, t(39)=8.157, p<0.0001, unpaired t-test. Muscle *draper* knockdown, t(47)=4.407, p<0.0001, unpaired t- test. Double knockdown, t(27)=8.579, p<0.0001, unpaired t-test. For B-E, N (NMJs) = 21, 26, 21, 20, 21, 28, 15, 14. Error bars indicate ± SEM, ns = non-significant, *p<0.05, **p<0.01, ***p<0.001, ****p<0.0001.

**Figure S5.**
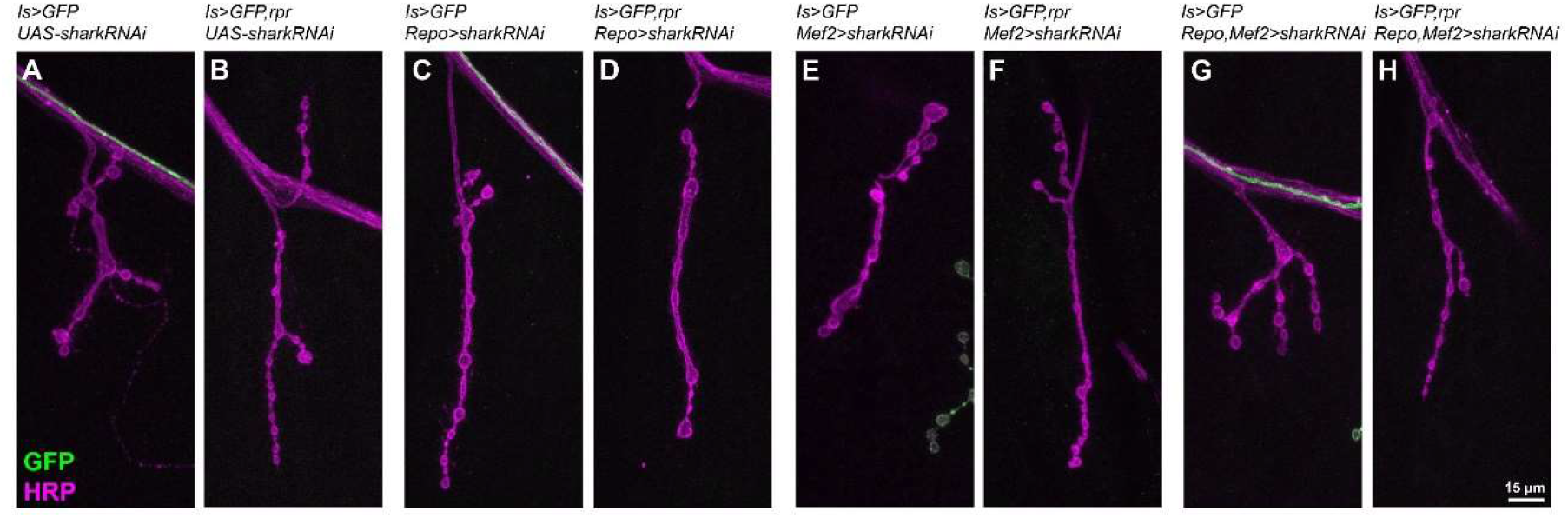
Representative NMJ images of cell type specific *shark* knockdown larvae. A-H. NMJs of MN4-Ib in third instar larvae of no ablation and Is ablated larvae in control, glia *shark* knockdown, muscle *shark* knockdown, and double knockdown backgrounds, labeled with GFP (green) and HRP (magenta). NMJ expansion was observed upon Is MN ablation (B), which is blocked by glia *shark* knockdown (D) or double knockdown (H).

**Figure S6.**
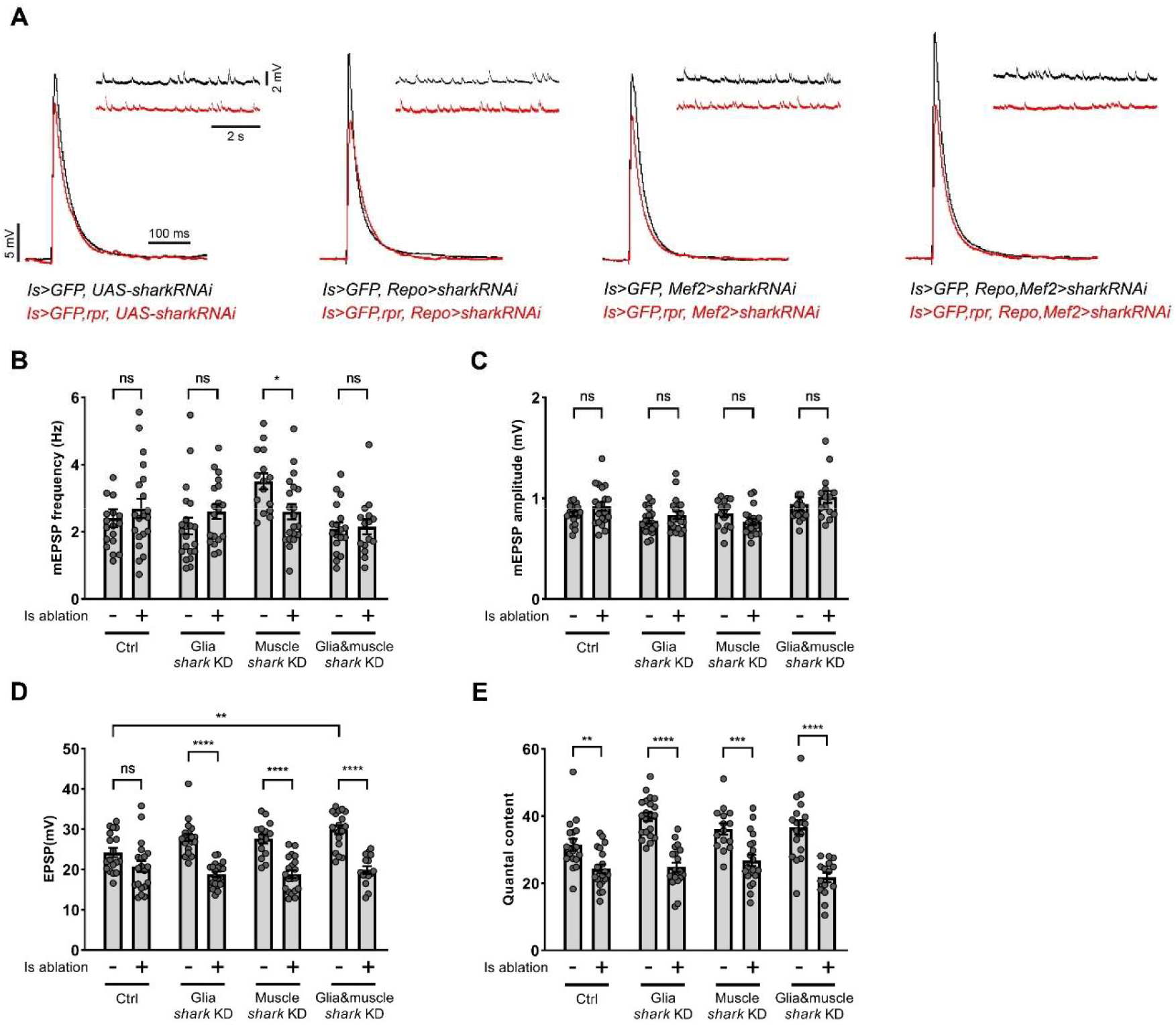
Electrophysiology recordings in cell type specific *shark* knock down larvae. A. EPSP and mEPSP recordings from no ablation and Is ablated larvae in control, glia *shark* knockdown, muscle *draper* knockdown and double knockdown backgrounds. B. Quantification of mEPSP frequency of no ablation and Is ablated larvae in control, glia *shark* knockdown, muscle *shark* knockdown, and double knockdown backgrounds. Control, t(36)=0.7421, p=0.4629, unpaired t-test. Glia *draper* knockdown, t(39)=1.300, p=0.2015, unpaired t-test. Muscle *draper* knockdown, t(33)=2.710, p=0.0106, unpaired t-test. Double knockdown, t(31)=0.1874, p=0.8525, unpaired t-test. C. Quantification of mEPSP amplitude of no ablation and Is ablated larvae in control, glia *shark* knockdown, muscle *shark* knockdown, and double knockdown backgrounds. Control, t(29.11)=1.575, p=0.4629, unpaired t-test with Welch’s correction. Glia *draper* knockdown, t(39)=1.261, p=0.2149, unpaired t-test. Muscle *draper* knockdown, t(33)=1.748, p=0.0898, unpaired t-test. Double knockdown, t(31)=0.7594, p=0.4533, unpaired t-test. D. Quantification of EPSP amplitude of no ablation and Is ablated larvae in control, glia *shark* knockdown, muscle *shark* knockdown, and double knockdown backgrounds. Control, t(36)=1.889, p=0.0669, unpaired t-test. Glia *draper* knockdown, t(39)=8.230, p<0.0001, unpaired t-test. Muscle *draper* knockdown, t(33)=6.017, p<0.0001, unpaired t- test. Double knockdown, t(31)=6.823, p<0.0001, unpaired t-test. E. Quantification of quantal content of no ablation and Is ablated larvae in control, glia *shark* knockdown, muscle *shark* knockdown, and double knockdown backgrounds. Control, t(36)=3.328, p=0.0020, unpaired t-test. Glia *draper* knock down, t(39)=8.159, p<0.0001, unpaired t-test. Muscle *draper* knock down, t(33)=3.878, p=0.0005, unpaired t-test. Double knock down, t(31)=5.581, p<0.0001, unpaired t-test. For B-E, N (NMJs) = 18, 20, 21, 20, 15, 20, 18, 15. Error bars indicate ± SEM, ns = non-significant, *p<0.05, **p<0.01, ***p<0.001, ****p<0.0001.

**Figure S7.**
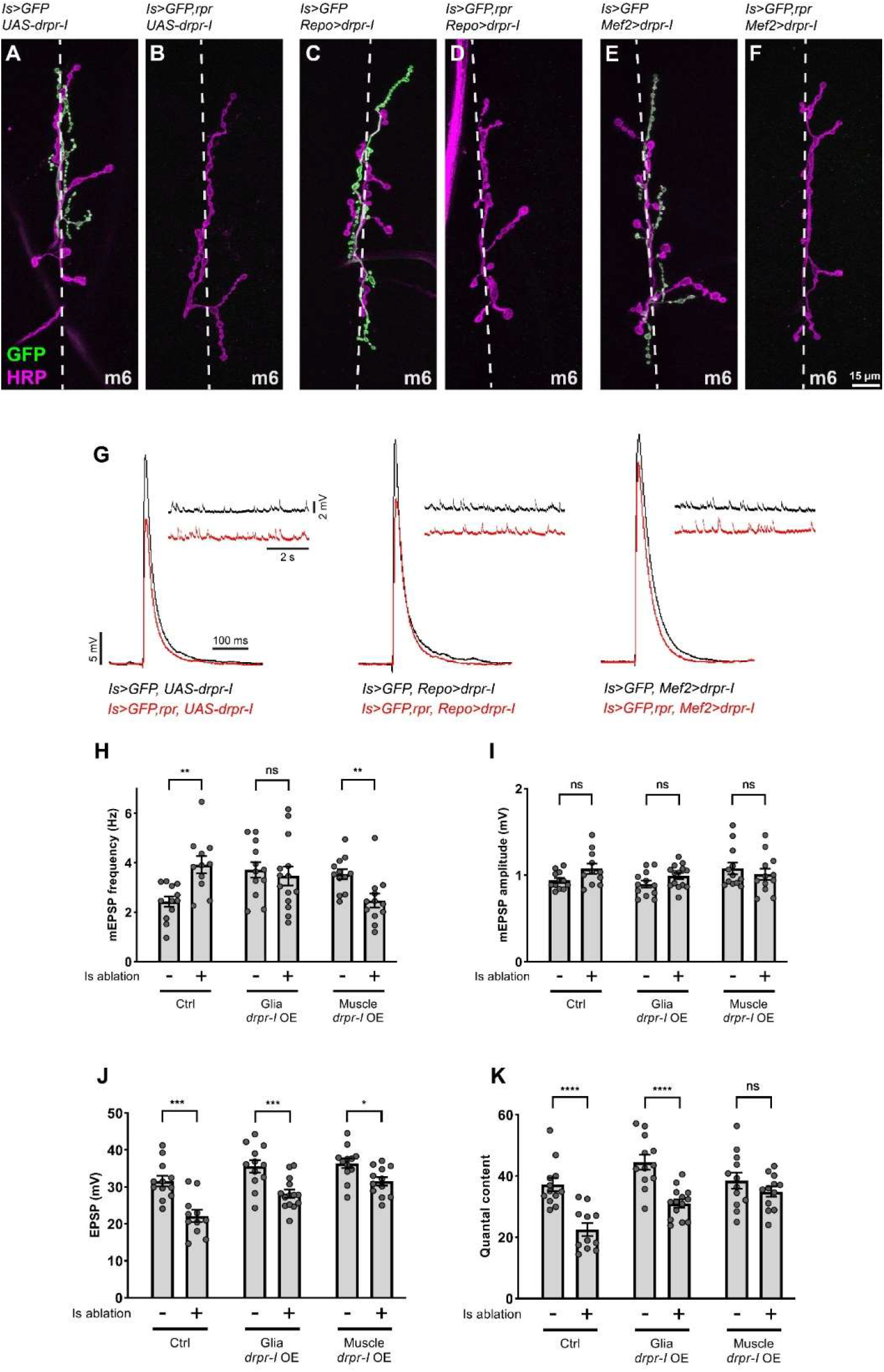
Morphology and physiology of MN6-Ib in *draper-I* overexpressing larvae. A-F. NMJs of MN6-Ib in third instar larvae of no ablation and Is ablated larvae in control, glia *draper-I* overexpression, and muscle *draper-I* overexpression backgrounds, labeled with GFP (green) and HRP (magenta). G. EPSP and mEPSP recordings from larvae of displayed genotypes. H. Quantification of mEPSP frequency of no ablation and Is ablated larvae in control, glia *draper-I* overexpression, and muscle *draper-I* overexpression backgrounds. Control, t(21)=3.734, p=0.0012, unpaired t-test. Glia *draper-I* overexpression, t(24)=0.4865, p=0.6310, unpaired t-test. Muscle *draper-I* overexpression, t(22)=3.059, p=0.0057, unpaired t-test. I. Quantification of mEPSP amplitude of no ablation and Is ablated larvae in control, glia *draper-I* overexpression, and muscle *draper-I* overexpression backgrounds. Control, t(14.48)=2.050, p=0.0589, unpaired t-test with Welch’s correction. Glia *draper-I* overexpression, t(24)=1.707, p=0.1007, unpaired t-test. Muscle *draper-I* overexpression, t(22)=0.7120, p=0.4839, unpaired t-test. J. Quantification of EPSP amplitude of no ablation and Is ablated larvae in control, glia *draper-I* overexpression, and muscle *draper-I* overexpression backgrounds. Control, t(21)=4.324, p=0.0003, unpaired t-test. Glia *draper-I* overexpression, t(24)=3.757, p=0.0010, unpaired t-test. Muscle *draper-I* overexpression, t(22)=2.806, p=0.0103, unpaired t-test. K. Quantification of quantal content of no ablation and Is ablated larvae in control, glia *draper-I* overexpression, and muscle *draper-I* overexpression backgrounds. Control, t(21)=4.858, p<0.0001, unpaired t-test. Glia *draper-I* overexpression, t(24)=5.024, p<0.0001, unpaired t-test. Muscle *draper-I* overexpression, t(22)=1.164, p=0.2569, unpaired t-test. For H-K, N (NMJs) = 12, 11, 12, 14, 12, 12. Error bars indicate ± SEM, ns = non-significant, *p<0.05, **p<0.01, ***p<0.001, ****p<0.0001.

**Figure S8.**
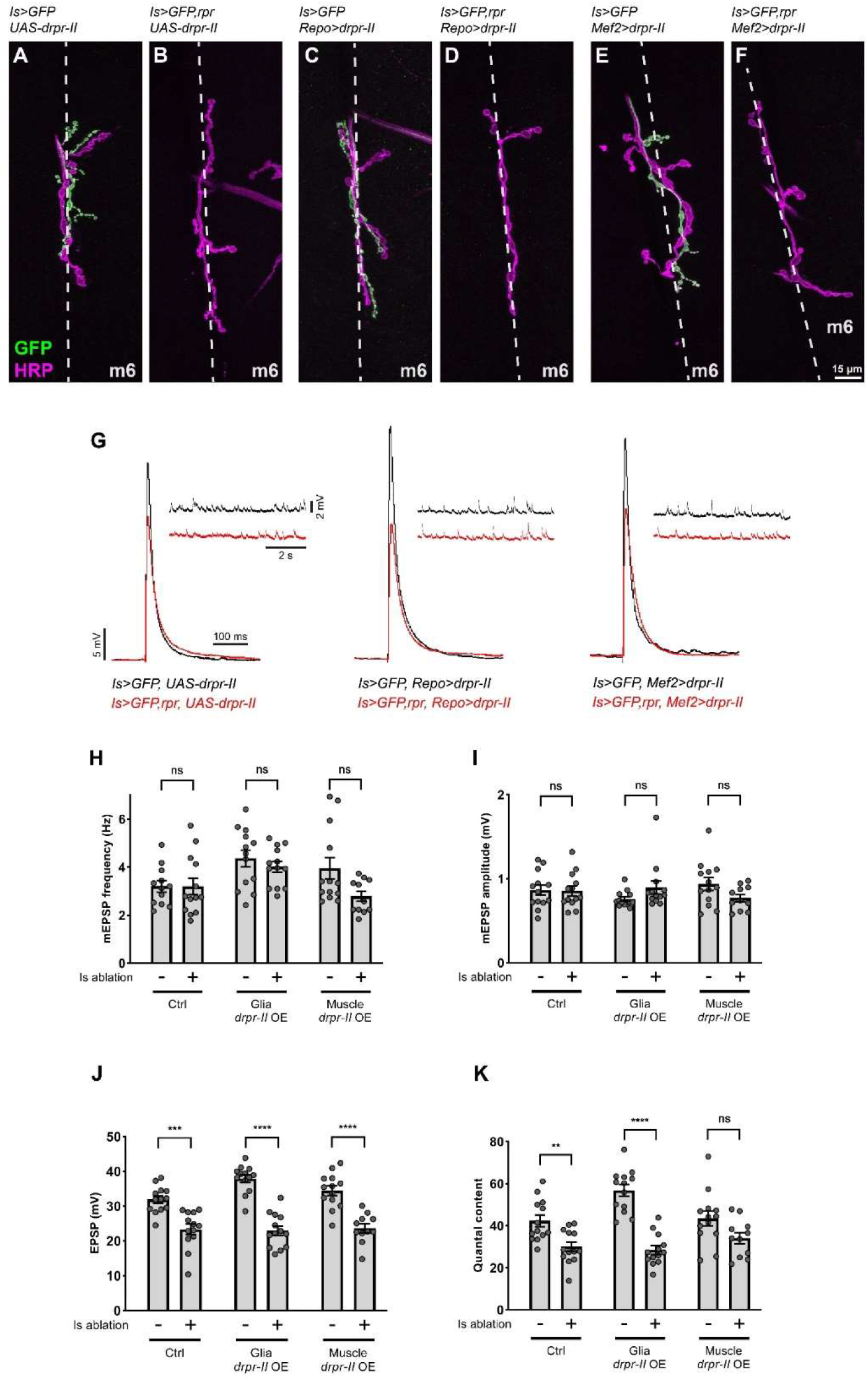
Morphology and physiology of MN6-Ib in *draper-II* overexpressing larvae. A-F. NMJs of MN6-Ib in third instar larvae of no ablation and Is ablated larvae in control, glia *draper-II* overexpression, and muscle *draper-II* overexpression background, labeled with GFP (green) and HRP (magenta). G. EPSP and mEPSP recordings from larvae of displayed genotypes. H. Quantification of mEPSP frequency of no ablation and Is ablated larvae in control, glia *draper-II* overexpression, and muscle *draper-II* overexpression backgrounds. Control, t(24)=0.0086, p=0.9932, unpaired t-test. Glia *draper-II* overexpression, t(24)=0.8540, p=0.4015, unpaired t-test. Muscle *draper-II* overexpression, t(22)=2.237, p=0.0358, unpaired t-test. I. Quantification of mEPSP amplitude of no ablation and Is ablated larvae in control, glia *draper-II* overexpression, and muscle *draper-II* overexpression backgrounds. Control, t(24)=0.0733, p=0.9422, unpaired t-test. Glia *draper-II* overexpression, t(24)=1.733, p=0.0959, unpaired t-test. Muscle *draper-II* overexpression, t(22)=1.841, p=0.0791, unpaired t-test. J. Quantification of EPSP amplitude of no ablation and Is ablated larvae in control, glia *draper-II* overexpression, and muscle *draper-II* overexpression backgrounds. Control, t(24)=4.647, p=0.0001, unpaired t-test. Glia *draper-II* overexpression, t(24)=8.613, p<0.0001, unpaired t-test. Muscle *draper-II* overexpression, t(22)=5.633, p<0.0001, unpaired t-test. K. Quantification of quantal content of no ablation and Is ablated larvae in control, glia *draper-II* overexpression, and muscle *draper-II* overexpression background. Control, t(24)=3.466, p=0.0020, unpaired t-test. Glia *draper-II* overexpression, t(24)=8.142, p<0.0001, unpaired t-test. Muscle *draper-II* overexpression, t(22)=2.044, p=0.0531, unpaired t-test. For H-K, N (NMJs) = 13, 13, 13, 13, 13, 11. Error bars indicate ± SEM, ns = non-significant, **p<0.01, ***p<0.001, ****p<0.0001.

**Figure S9.**
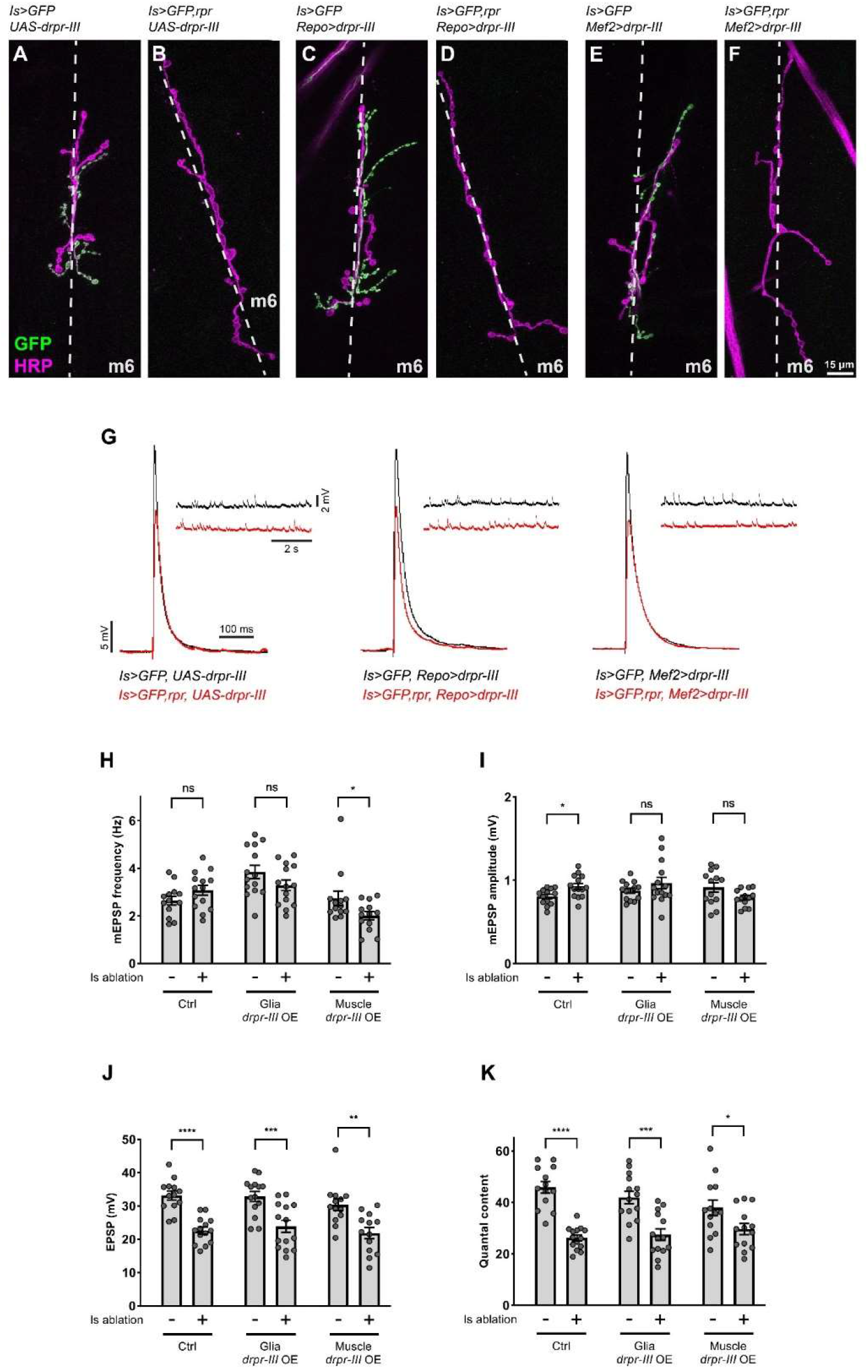
Morphology and physiology of MN6-Ib in *draper-III* overexpressing larvae. A-F. NMJs of MN6-Ib in third instar larvae of no ablation and Is ablated larvae in control, glia *draper-III* overexpression, and muscle *draper-III* overexpression backgrounds, labeled with GFP (green) and HRP (magenta). G. EPSP and mEPSP recordings from larvae of displayed genotypes. H. Quantification of mEPSP frequency of no ablation and Is ablated larvae in control, glia *draper-III* overexpression, and muscle *draper-III* overexpression backgrounds. Control, t(25)=1.544, p=0.1350, unpaired t-test. Glia *draper-III* overexpression, t(26)=1.541, p=0.1354, unpaired t-test. Muscle *draper-III* overexpression, t(24)=2.079, p=0.0485, unpaired t-test. I. Quantification of mEPSP amplitude of no ablation and Is ablated larvae in control, glia *draper-III* overexpression, and muscle *draper-III* overexpression backgrounds. Control, t(25)=2.727, p=0.0115, unpaired t-test. Glia *draper-III* overexpression, t(26)=1.224, p=0.2319, unpaired t-test. Muscle *draper-III* overexpression, t(24)=1.943, p=0.0638, unpaired t-test. J. Quantification of EPSP amplitude of no ablation and Is ablated larvae in control, glia *draper-III* overexpression, and muscle *draper-III* overexpression backgrounds. Control, t(25)=6.298, p<0.0001, unpaired t-test. Glia *draper-III* overexpression, t(26)=3.814, p=0.0008, unpaired t-test. Muscle *draper-III* overexpression, t(24)=3.549, p=0.0016, unpaired t-test K. Quantification of quantal content of no ablation and Is ablated larvae in control, glia *draper-III* overexpression, and muscle *draper-III* overexpression backgrounds. Control, t(25)=7.938, p<0.0001, unpaired t-test. Glia *draper-III* overexpression, t(26)=4.425, p=0.0002, unpaired t-test. Muscle *draper-III* overexpression, t(24)=2.208, p=0.0371, unpaired t-test. For H-K, N (NMJs) = 13, 14, 14, 14, 13, 13. Error bars indicate ± SEM, ns = non-significant, *p<0.05, **p<0.01, ***p<0.001, ****p<0.0001.

**Figure S10.**
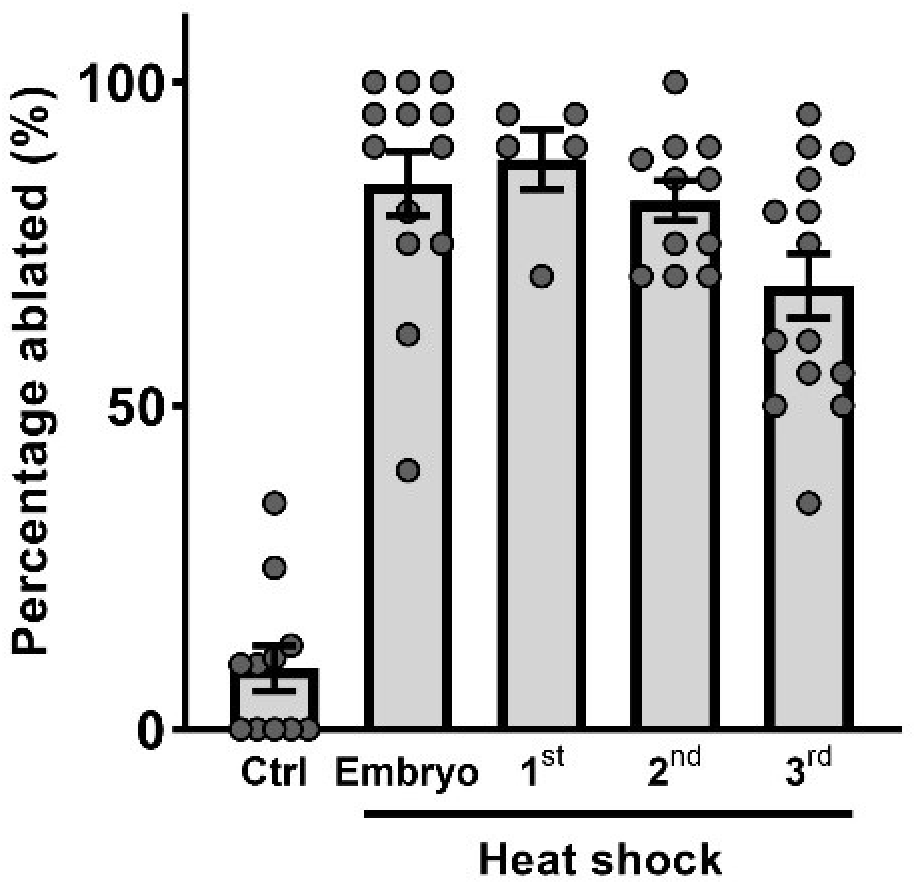
Examining the ablation efficiency of the heat-shock induced Is MN ablation system. Heat-shock at different developmental stages (embryo, first, second, and third instar) induces a high percentage of Is MN ablation.

**Figure S11.**
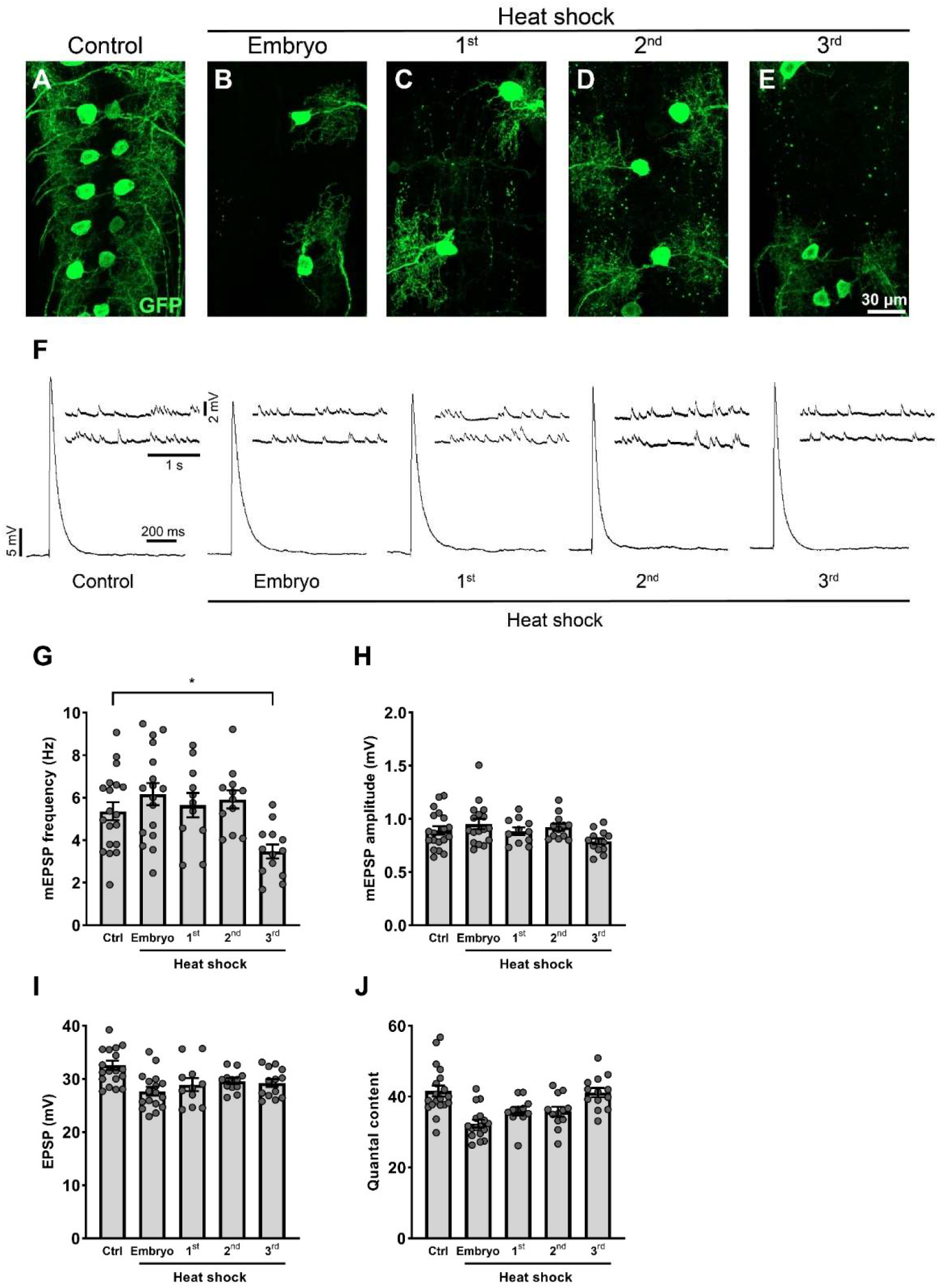
Morphology and electrophysiology of larvae with Is MNs ablated at different developmental stages. A-E. VNCs of late third instar larvae (*hs-FLP,UAS-GFP/+;UAS-FRT stop FRT-hid-2A- rpr/+;Is-GAL4/+*) with (A) no heat-shock, (B) embryo heat-shock, (C) first instar heat-shock, (D) second instar heat-shock, and (E) third instar heat-shock, stained with GFP (green). The Is MN cell bodies are fully removed in all conditions. F. EPSP and mEPSP recordings from larvae with Is MNs ablated at different developmental stages. G. Quantification of mEPSP frequency. F(4, 67)=4.926, p=0.0015. Control vs third instar heat-shocked, p=0.0340. H. Quantification of mEPSP amplitude. F(4, 67)=2.248, p=0.0730. I. Quantification of EPSP amplitude. F(4, 67)=5.399, p=0.0008. J. Quantification of quantal content. F(4, 67)=9.109, p<0.0001. For G-J, N (NMJs) = 19, 17, 11, 12, 13. Error bars indicate ± SEM, *p<0.05.

**Video 1. Roll behaviors of control and Is ablated larvae.**

